# RUNX1 is required in granulocyte-monocyte progenitors to attenuate inflammatory cytokine production by neutrophils

**DOI:** 10.1101/2023.01.27.525911

**Authors:** Alexandra U. Zezulin, Darwin Ye, Elizabeth Howell, Daniel Yen, Erica Bresciani, Jamie Diemer, Jian-gang Ren, Mohd Hafiz Ahmad, Lucio H. Castilla, Ivo P. Touw, Andy J. Minn, Wei Tong, P. Paul Liu, Kai Tan, Wenbao Yu, Nancy A. Speck

## Abstract

The transcription factor RUNX1 is mutated in familial platelet disorder with associated myeloid malignancies (FPDMM) and in sporadic myelodysplastic syndrome and leukemia. RUNX1 regulates inflammation in multiple cell types. Here we show that RUNX1 is required in granulocyte-monocyte progenitors (GMPs) to restrict the inflammatory response of neutrophils to toll-like receptor 4 (TLR4) signaling. Loss of RUNX1 in GMPs increased the TLR4 coreceptor CD14 on neutrophils, which contributed to neutrophils’ increased inflammatory cytokine production in response to the TLR4 ligand lipopolysaccharide. RUNX1 loss increased the chromatin accessibility of retrotransposons in GMPs and neutrophils and induced a type I interferon signature characterized by enriched footprints for signal transducer and activator of transcription (STAT1::STAT2) and interferon regulatory factors (IRF) in opened chromatin, and increased expression of interferon-stimulated genes. The overproduction of inflammatory cytokines by neutrophils was reversed by inhibitors of type I IFN signaling. We conclude that RUNX1 restrains the chromatin accessibility of retrotransposons in GMPs and neutrophils, and that loss of RUNX1 increases proinflammatory cytokine production by elevating tonic type I interferon signaling.

## INTRODUCTION

The transcription factor (TF) RUNX1 is mutated in both sporadic and inherited forms of leukemia and myelodysplastic syndrome (MDS) ^1^. Inherited *RUNX1* mutations cause familial platelet disorder with associated myeloid malignancy (FPDMM) which is characterized by thrombocytopenia, platelet activation defects, accelerated clonal hematopoiesis, and an up to 50% lifetime risk of leukemia ^2-4^. The inflammatory disease eczema was reported in an FPDMM pedigree ^5^, and since then has been documented in up to 50% of families ^2,3^. FPDMM patients also have an increased incidence of asthma, reactive airway disease, rosacea, and allergies ^6^. Chronic inflammation is a driver of leukemia ^7,8^, therefore inflammatory disorders in FPDMM patients may contribute to the elevated incidence of hematologic malignancies ^1^.

The regulation of inflammation by RUNX1 has been observed in multiple experimental contexts and occurs via diverse mechanisms. RUNX1 has been shown to promote, but more often to repress inflammation. RUNX1 promotes inflammatory signaling in macrophages by binding the p50 subunit of nuclear factor kappa B (NF-kB), which augments the expression of several pro-inflammatory cytokine genes in response to toll-like receptor 4 (TLR4) signaling ^9^. RUNX1 represses inflammation through its interactions with FOXP3, an essential transcription factor in regulatory T cells (Tregs) ^10^. RUNX1 represses inflammation in non-immune lung alveolar epithelial cells in response to lipopolysaccharide (LPS)-induced lung injury by inhibiting IkB kinase ^11^.

RUNX1 also represses inflammation by negatively regulating type I interferon (IFN) production and signaling. Type I IFN signaling is triggered by binding of type I IFN to the transmembrane IFNα/β receptor. The activated receptor phosphorylates and activates Janus activated kinase 1 (JAK1) and tyrosine kinase 2 (TYK2), which in turn phosphorylate the signal transducer and activator of transcription (STAT) proteins STAT1, and STAT2. Phosphorylated STAT1 (p-STAT1), p-STAT2, and an IFN regulatory factor assemble into a complex that translocates into the nucleus and activates the transcription of a group of IFN-stimulated genes (ISGs). Overexpression of dominant negative forms of RUNX1, or RUNX1 knockdown induced the expression of type I IFNs and ISGs, while overexpressing RUNX1 had the opposite effect ^12,13^. Deletion of RUNX1 in B cells also increased the expression of ISGs and caused them to hyper - respond to LPS following pre-stimulation of the B cell receptor ^14^.

Type I IFN production can be triggered by the de-repression of transposable elements (TEs). TEs consist of retrotransposons and DNA transposons, which together constitute a large percentage of the mammalian genome. Retrotransposons can be divided into two groups—the long terminal repeats (LTR) and endogenous retroviruses (LTR/ERVs) and non-LTR elements including long or short interspersed nuclear elements (LINE and SINE). The chromatin accessibility of TEs is held in check by multiple epigenetic mechanisms including DNA methylation and histone modification, but in certain disease states TEs can become de-repressed ^15^. LTR/ERVs, and in particular ERVK, often escape epigenetic silencing in both hematologic and solid malignancies ^16,17^, and dysregulated retrotransposon expression has been linked to inflammation in autoimmune and neurological diseases ^18^. LTR/ERVs contain STAT1 binding sites and act as enhancers for ISGs ^19^. In addition, bidirectional transcription of retrotransposons leads to the production of dsRNA which can trigger the innate immune response by binding cytosolic RIG-I-like receptors which activate a signaling pathway that culminates in the expression of type I IFNs ^20^. A study of 178 AML patients from the Cancer Genome Atlas (TCGA) found that patients with RUNX1 mutations had altered levels of transcripts from TEs, suggesting a role for RUNX1 in regulating their expression ^21^.

We previously showed that pan-hematopoietic RUNX1 deletion resulted in bone marrow (BM) neutrophils that secreted elevated amounts of tumor necrosis factor (TNF), macrophage inhibitory protein α (MIP-1α, or CCL3), and IL-1α following activation of TLR4 signaling ^22^. However, deleting RUNX1 in more differentiated neutrophils using a neutrophil-specific Cre (S100A8-Cre) did not affect the production of inflammatory molecules, indicating that RUNX1 protein was not functioning in neutrophils *per se* to regulate TLR4 signaling ^22^. We hypothesized that instead RUNX1 was required in a neutrophil precursor to restrain inflammatory cytokine production by neutrophils. Single cell RNA sequence analysis of RUNX1 deficient hematopoietic progenitors hinted that the dysregulation of inflammatory signaling occurred at the granulocyte-monocyte progenitor (GMP) stage ^22^. Here we show RUNX1 is required in GMPs to restrain inflammatory cytokine production by neutrophils. Loss of RUNX1 in GMPs upregulates an inflammatory transcriptional program in both GMPs and neutrophils, characterized by an elevated type I IFN signature. Epigenetic profiling by Assay for Transposase-Accessible Chromatin with high throughput sequencing (ATAC-seq) and digital footprinting determined that the chromatin associated with both protein-coding genes and retrotransposons that gained accessibility upon RUNX1 loss was highly enriched for footprints for STAT1::STAT2 and multiple IRFs, providing further evidence that RUNX1 regulates type I IFN signaling in neutrophils and GMPs. Treatment of neutrophils with inhibitors of type I IFN signaling ameliorated the hyper-response of RUNX1 deficient neutrophils to LPS. Together these data indicate that the hyper-response of neutrophils to LPS-induced TLR4 signaling is a consequence of RUNX1’s absence in GMPs which elevates tonic type I IFN signaling in both GMPs and neutrophils.

## RESULTS

### RUNX1 is required in GMPs to restrain inflammatory cytokine production by neutrophils

We used several complementary approaches to determine if the hyper-response to LPS was an intrinsic property of RUNX1 deficient neutrophils, or an indirect effect of RUNX1 loss in another hematopoietic cell lineage. First, we addressed whether the aberrant neutrophil response was secondary to RUNX1 loss in lymphocytes (for example loss of Tregs) by comparing the LPS response of neutrophils from mice in which RUNX1 was deleted in all hematopoietic cells by Vav1-Cre (hereafter referred to as Runx1^ΔHSC^), to neutrophils from mice in which *Runx1* was deleted only in lymphoid cells by Rag1-Cre (Runx1^ΔLym^) (Fig. 1A). We stimulated BM cells from Runx1^ΔHSC^, Runx1^ΔLym^, or Control (*Runx1*^*f/f*^) mice *ex vivo* with vehicle or LPS for 4 hours, then measured the percentage of TNF^+^ neutrophils by intracellular flow cytometry (see Fig. S1A for gating strategy). The percentage of neutrophils that were TNF^+^ following LPS treatment was greater in Runx1^ΔHSC^ compared to Control BM cells, but not in Runx1^ΔLym^ BM, indicating that the loss of RUNX1 in lymphoid cells alone does not cause neutrophils to hyper-respond to LPS (Fig. 1B, S1B). Since lymphocytes were not completely absent in Runx1^ΔLym^ mice (Fig. S1C,D), we also examined the inflammatory response of neutrophils from *Rag2*^*-/-*^ mice which lack lymphocytes ^23^ (Fig. S1E,F). Purified neutrophils from *Rag2*^*-/-*^ mice did not overproduce either TNF or CCL3, as determined using a cytometric bead array (CBA) (Fig. 1C,D), confirming that neutrophils from Runx1^ΔHSC^ mice did not hyper-respond because of lymphocyte defects. To address whether other hematopoietic cells in the BM could be responsible for the hyper-responsive neutrophil phenotype, we generated BM chimeras by transplanting a 10:1 ratio of BM cells from Control (C57BL6/J x B6.SJL) F1 and Runx1^ΔHSC^ mice into irradiated B6.SJL mice (Fig. 1E). At 24 weeks post-transplant, 100% of T and B cells and 97% of the neutrophils in recipient mice were derived from the transplanted Control cells, thus they far outnumber the Runx1^ΔHSC^ neutrophils (Fig. 1F). Runx1^ΔHSC^ neutrophils purified from transplant recipient mice secreted more TNF and CCL3 than Control neutrophils purified from the same recipient mice (Fig. 1G). Therefore, Runx1^ΔHSC^ neutrophils in a BM consisting primarily of normal hematopoietic cells hyper-respond to LPS stimulation, consistent with a neutrophil-intrinsic defect.

**Figure 1.**
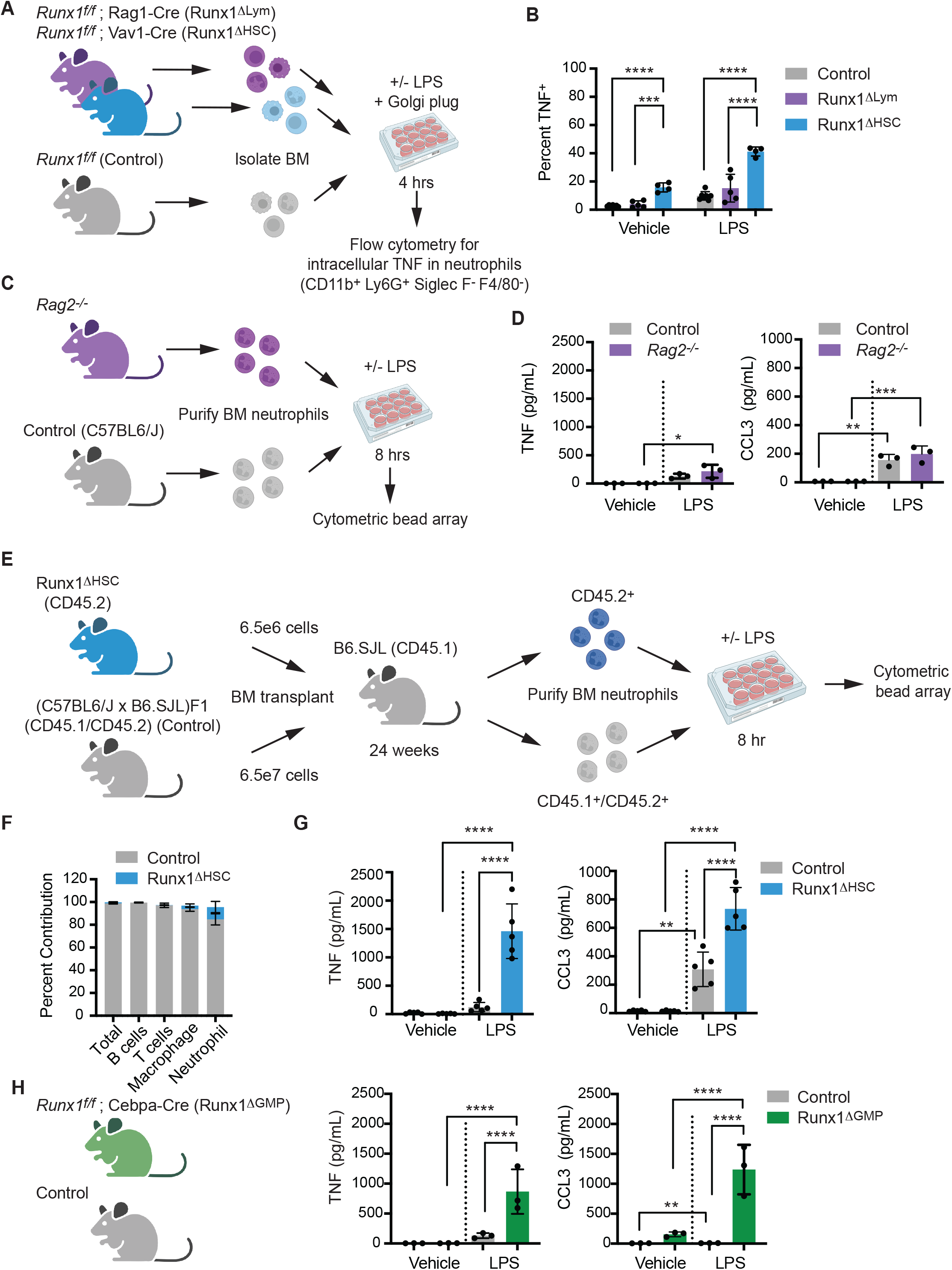
RUNX1 function in GMPs is necessary to restrict inflammatory cytokine production by neutrophils. **A)** Schematic depicting experiment to compare the effect of pan-hematopoietic (Runx1^ΔHSC^) versus lymphocyte-specific (Runx1^ΔLym^) RUNX1 loss on inflammatory cytokine production by neutrophils. BM cells were stimulated for 4 hours *ex vivo* with vehicle or 100 ng/mL LPS for 4 hours. The percentage of neutrophils that were TNF^+^ was determined by flow cytometry. **B)** Percentage of Runx1^ΔHSC^, Runx1^ΔLym^, and Control BM neutrophils that were TNF^+^. Mean ± SD, one way ANOVA, Tukey’s multiple comparison test, representative of 3 experiments. **C)** Schematic depicting experiment to examine the inflammatory phenotype of neutrophils from *Rag2*^*-/-*^ mice. **D)** Absolute quantification by CBA of inflammatory factors in the supernatant of neutrophils from Control and *Rag2*^*-/-*^ mice stimulated for 8 hours with vehicle or 100 ng/mL LPS. Mean ± SD, one-way ANOVA plus Tukey multiple comparison test. Representative of 2 experiments. **E)** Schematic representation of experimental design for generating BM chimeras by transplanting Runx1^ΔHSC^ BM cells and a 10-fold excess of Control BM cells into irradiated B6.SJL mice. Neutrophils derived from transplanted Runx1^ΔHSC^ and Control BM were purified by FACS 24 weeks post-transplant, and analyzed as in panels C,D. **F)** Percentage of total BM cells, B cells (CD19^+^), T cells (CD3^+^), granulocytes (Gr1^+^CD11b^+^), and macrophages (CD11b^+^Gr1^-^) derived from Runx1^ΔHSC^ versus Control BM in transplant recipient mice. **G)** Representative experiment of absolute quantification by CBA of inflammatory factors in the supernatant of Control and Runx1^ΔHSC^ neutrophils purified from transplant recipients and stimulated for 8 hours with vehicle or 100 ng/mL LPS. Mean ± SD, one-way ANOVA plus Tukey multiple comparison test, representative of 2 experiments. **H)** Absolute quantification by CBA of inflammatory factors in the supernatant of BM neutrophils from Control or Runx1^ΔGMP^ mice, analyzed as described in panel C. Mean ± SD, one-way ANOVA plus Tukey multiple comparison test, representative of 9 experiments. For all experiments; *P≤0.05, **P≤0.01 ***P≤0.001, ****P ≤ 0.0001.

In our previous scRNA-seq analysis of Runx1^ΔHSC^ hematopoietic stem and progenitor cells, an aberrant inflammatory transcriptional signature was first detectable in GMPs ^22^. To test whether RUNX1 loss in GMPs was sufficient to generate hyper-responsive neutrophils, we deleted RUNX1 using Cebpa-Cre (Runx1^ΔGMP^), which deletes primarily in GMPs (76%), and to a lesser extent in common lymphoid progenitors (9%), common myeloid progenitors (26%) and LSK cells (13%) ^24^. Cebpa-Cre efficiently deleted RUNX1 in GMPs and neutrophils (Fig. S1G). BM neutrophils from Runx1^ΔGMP^ mice overproduced TNF and CCL3 in response to LPS (Fig. 1H), demonstrating that loss of RUNX1 in GMPs is sufficient to establish a hyper-responsive neutrophil phenotype.

### RUNX1 loss results in elevated levels of key TLR4 signaling molecules

We analyzed the basal transcriptional changes (in the absence of LPS) in GMPs and neutrophils purified from Runx1^ΔGMP^ mice by bulk RNA-seq (data quality are documented in Fig. S2A,B). ∼1.2K genes were differentially expressed in Control versus Runx1^ΔGMP^ GMPs, and 954 in Control versus Runx1^ΔGMP^ neutrophils, with more genes downregulated than upregulated in each cell type (Fig. 2A). Multiple enriched Gene Ontology (GO) terms for genes upregulated in Runx1^ΔGMP^ GMPs and neutrophils were related to inflammatory and immune responses, including “response to interferon beta”, “defense response to protozoan”, and “neutrophil extracellular trap formation” (Fig. 2B). The data indicate that RUNX1 loss de-represses an immune response transcriptional program in both GMPs and neutrophils.

**Figure 2.**
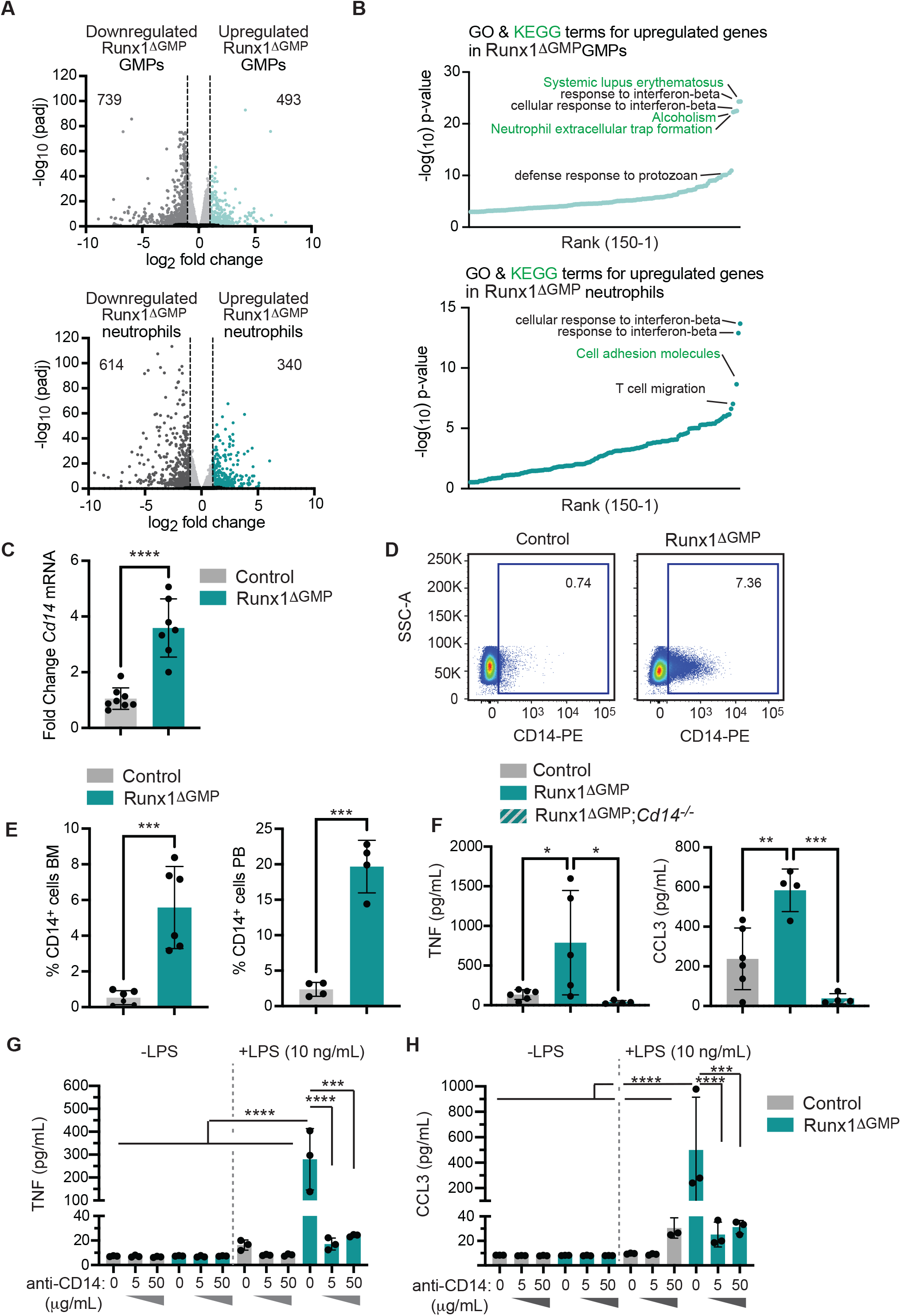
RUNX1 loss results in elevated levels of key TLR4 signaling molecules. **A)** Volcano plots depicting global transcriptional changes between Control and Runx1^ΔGMP^ GMPs and neutrophils. Upregulated genes with an adjusted p-value less than 0.05 and a log_2_ fold change greater than 1 are indicated by teal dots, and downregulated genes with a log_2_ fold change less than −1 by dark gray dots. The numbers of significantly up or downregulated genes are indicated. **B)** Top 150 enriched Gene ontology (GO) (black text) and Kyoto Encyclopedia of Genes and Genomes (KEGG) pathway terms (green text) for genes upregulated in Runx1^ΔGMP^ GMPs and neutrophils. The enrichment of GO and KEGG terms was tested using Fisher exact test (GeneSCF v1.1-p2). **C)** RT-qPCR for *Cd14* in Runx1^ΔGMP^ neutrophils. Mean ± SD, unpaired, two-tailed Student’s t test. **D)** Representative scatter plots of CD14 expression on BM neutrophils from Control and Runx1^ΔGMP^ mice. **E)** Quantification of the percentage of CD14^+^ neutrophils in the BM (left) and PB (right) of Control and Runx1^ΔGMP^ mice. Mean ± SD, unpaired, two-tailed Student’s t test, representative of 11 experiments, a total of 32 mice were analyzed. **F)** Absolute quantification by CBA demonstrating deletion of *Cd14* reduces TNF and CCL3 production by Control and Runx1^ΔGMP^ neutrophils in response to 10 ng/mL LPS. Mean ± SD, one-way ANOVA plus Tukey multiple comparison test, representative of 2 experiments, a total of 24 mice were analyzed. **G)** Absolute quantification by CBA demonstrating effect of CD14 blocking antibody (50 µg/mL) on TNF production by purified BM-derived neutrophils stimulated for 8 hours with vehicle or a low dose (10 ng/mL) of LPS. Mean ± SD, one-way ANOVA plus Tukey multiple comparison test, representative of 2 experiments, a total of 8 mice were analyzed. **H)** Effect of CD14 blocking antibody on CCL3 production by purified neutrophils stimulated for 8 hours with vehicle or a low dose of LPS, as in panel G. For all figures; P≤0.05, **P≤0.01, ***P=0.001, ****P ≤ 0.0001.

Expression of the *Cd14* gene was upregulated in Runx1^ΔGMP^ neutrophils by RNA-seq. CD14 is a glycosylphosphatidylinositol-anchored TLR4 accessory protein present on the cell membrane and secreted in a soluble form, which binds LPS and transfers it to TLR4 ^25^. The transfer of LPS from CD14 to TLR4 facilitates TLR4 endocytosis, activation of the TRIF pathway, and type I IFN production ^25,26^. CD14 mRNA was elevated approximately 2-fold by RT-qPCR (Fig. 2C), and the percentages of CD14^+^ neutrophils in the BM and peripheral blood (PB) were increased by ∼10 fold (Fig. 2D,E, Fig. S3A). CD14 contributed to the hyper-response, as neutrophils from Runx1^ΔGMP^ mice deficient in CD14 ^27^ produced lower levels of TNF and CCL3 in response to LPS (Fig. 2F). CD14 increases the sensitivity of TLR4 activation under conditions of low (picomolar) LPS concentrations but is not required for TLR4 responses to high LPS concentrations ^25,28^. Blocking antibodies against CD14 decreased the levels of TNF and CCL3 that were secreted into culture supernatants by Runx1^ΔGMP^ neutrophils in response to a low concentration of LPS (10 ng/mL) (Fig. 2G,H) but not to a 10-fold higher concentration (100 ng/mL) (Fig. S3B). Therefore, elevated levels of CD14 contribute to the hyper-response of Runx1^ΔGMP^ neutrophils to low levels of LPS, and inhibiting CD14 could dampen the exaggerated response.

### Loss of RUNX1 in GMPs increases the chromatin accessibility of genes involved in innate immune responses in both GMPs and neutrophils

We hypothesized that epigenetic alterations in key inflammatory pathway genes are acquired in Runx1^ΔGMP^ GMPs and propagated to neutrophils. To examine this, we performed Assay for Transposon-Accessible Chromatin (ATAC)-seq on non-LPS-stimulated GMPs and neutrophils. Overall, despite the fact that fewer genes were expressed in Runx1^ΔGMP^ GMPs and neutrophils, more ATAC-seq peaks were gained than lost (Figs. 3A,B). Of the 5579 peaks that were higher in Runx1^ΔGMP^ neutrophils relative to Control neutrophils, 75% had been strongly (2442) or mildly (1736) gained in Runx1^ΔGMP^ GMPs (Fig. 3C). Therefore, most of the increases in chromatin accessibility in Runx1^ΔGMP^ neutrophils originated in Runx1^ΔGMP^ GMPs. In contrast, only 36% of peaks lost in Runx1^ΔGMP^ neutrophils were also lost in GMPs (Fig. S4A). Many GO terms specifically enriched for peaks that were lost in Runx1^ΔGMP^ GMPs or neutrophils (i.e. associated with peaks higher in Control GMPs and neutrophils) are related to developmental or cell biological processes (Fig. 3D,E). On the other hand, terms associated with peaks gained in Runx1^ΔGMP^ GMPs or neutrophils (Fig. 3F,G), or gained peaks shared by Runx1^ΔGMP^ GMPs and neutrophils (Fig. S4B, Table S2) were related to immune responses, suggesting that RUNX1 loss increased the accessibility of genes associated with immune cell activation beginning at the GMP stage. *Tlr4* and *Cd14* were included among the innate immune response genes with increased chromatin accessibility in both Runx1^ΔGMP^ GMPs and neutrophils (Fig. 3H). Peaks from ChIP-seq for the active enhancer mark H3K27ac also generated GO terms related to innate immune responses in Runx1^ΔGMP^ neutrophils (Fig. S4C,D), confirming that the enhancers of these genes were in a more active state.

**Figure 3.**
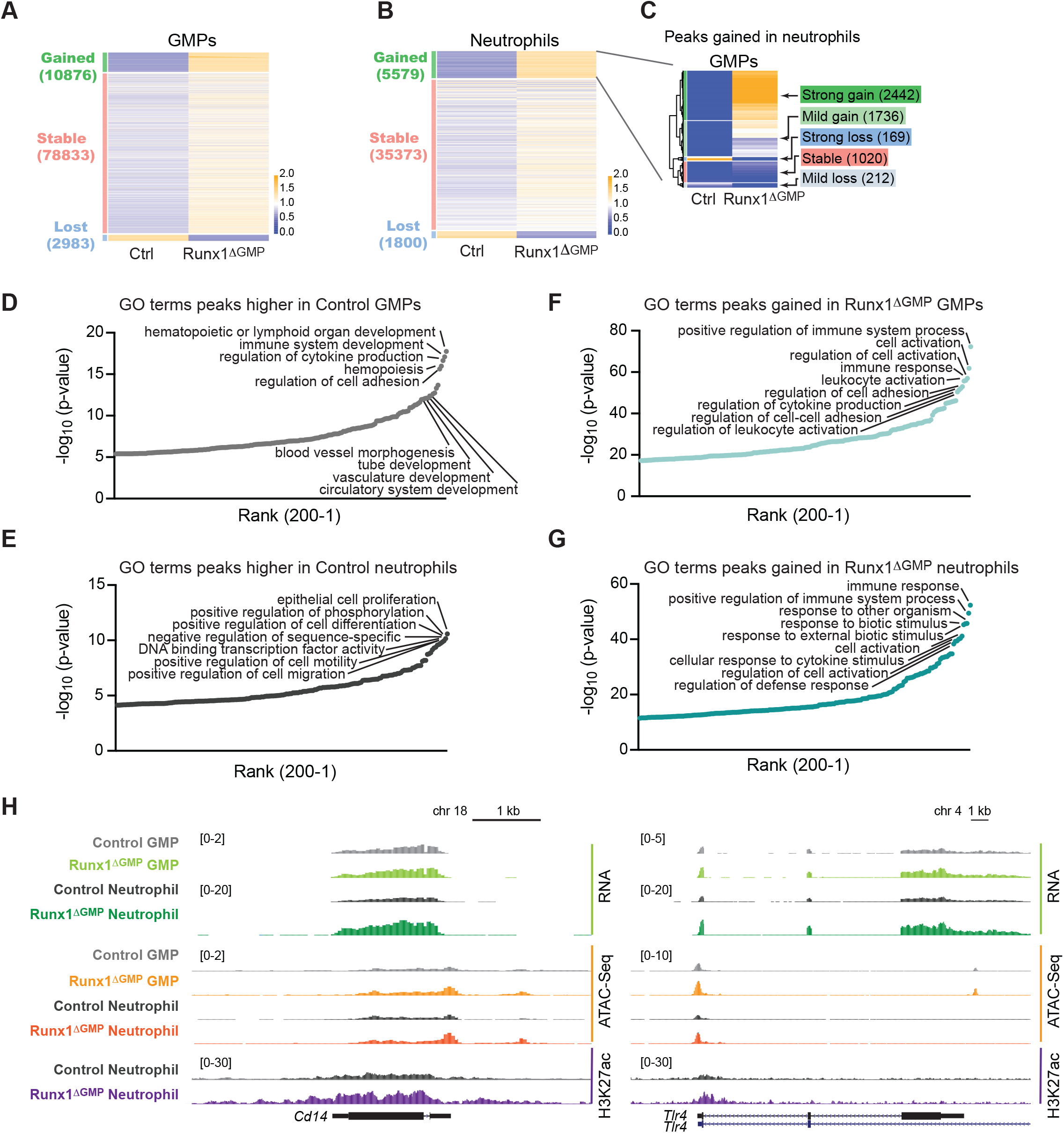
Loss of RUNX1 increases the chromatin accessibility of genes involved in innate immune responses in GMPs and neutrophils. **A)** Heatmap of ATAC-seq signals for Control (Ctrl) and Runx1^ΔGMP^ GMPs. Peaks are categorized as Gained, Lost (in Runx1^ΔGMP^ GMPs), or Stable (RPKM fold change <2).The color represents the relative RPKM to the mean RPKM of Control and Runx1^ΔGMP^ cells. **B)** Heatmap of ATAC-seq signals in Control and Runx1^ΔGMP^ neutrophils. **C)** Heatmap of ATAC-seq signal in Control and Runx1^ΔGMP^ GMPs for peaks gained in Runx1^ΔGMP^ neutrophils. The peaks were clustered hierarchically and then segregated into 5 groups using the cutree function in R. **D)** Enriched GO Biological terms for peaks higher in Control GMPs (i.e., peaks lost in Runx1^ΔGMP^ GMPs). The top 200 GO Biological terms are plotted. **E)** Enriched GO Biological terms for peaks higher in Control neutrophils (i.e., peaks lost in Runx1^ΔGMP^ neutrophils). **F)** Enriched GO Biological terms for peaks gained in Runx1^ΔGMP^ GMPs. **G)** Enriched GO Biological terms for peaks gained in Runx1^ΔGMP^ neutrophils. **H)** Genome browser view showing normalized RNA-seq, ATAC-seq, and H3K27ac signals for the *Tlr4* and *Cd14* genes in Control and Runx1^ΔGMP^ GMPs and neutrophils.

In conclusion, the chromatin of genes related to innate immune responses is more accessible in Runx1^ΔGMP^ GMPs, and this increased accessibility is propagated to Runx1^ΔGMP^ neutrophils.

### RUNX1 restrains tonic type I IFN signaling

We inferred where RUNX1 binds to chromatin in GMPs and neutrophils by digital footprinting ^29^. Digital ‘footprints’ in ATAC-seq data result from the Tn5 enzyme’s inability to cleave where TFs are bound to DNA, which results in a dip in reads, or ‘footprint’ in peaks that can then be matched to TF motifs to infer the specific TF bound. We identified ∼10K RUNX1 footprints in GMPs, and ∼3.5K in neutrophils (Table S3). We inferred the ability of RUNX1 to modify the accessibility of chromatin at its bound sites by determining differences in the average normalized read counts in 200 bp of sequence flanking the footprints ^29^. There was a trend towards decreased accessibility of chromatin surrounding the RUNX1 footprints in both GMPs and neutrophils when RUNX1 was lost (Fig. S5A), suggesting that RUNX1’s predominant activity is to open flanking chromatin at both stages. We next determined whether the binding of other TFs was affected at sites that lost or gained accessibility in Runx1^ΔGMP^ cells using the algorithm bias-free transcription factor Footprint Enrichment Test (biFET) ^30^. Footprints of all three RUNX TFs and multiple GATA TFs were highly enriched in peaks that were lost in GMPs when RUNX1 was deleted (Fig. S5B and Table S3). Footprints for RUNX TFs were also highly enriched in regions of decreased accessibility in Runx1^ΔGMP^ neutrophils, as were footprints for several other TFs (e.g., Bhlha15, FOXB1, ZSCAN4), although they were less enriched than RUNX footprints (Fig. S5B).

The most highly enriched TF footprints in ATAC-seq peaks gained in both Runx1^ΔGMP^ GMPs and neutrophils were for multiple IRFs and STAT1::STAT2 (Fig. 4A). Footprints for several E-twenty-six (ETS) TFs, SPI1 (PU.1), SPIB, and SPIC were also enriched in Runx1^ΔGMP^ neutrophils (Fig. 4A). The chromatin surrounding the IRF2, IRF3, IRF4, IRF9, STAT1::STAT2, and SPIC footprints was significantly more accessible in Runx1^ΔGMP^ neutrophils (Fig. 4B, Table S3), indicating that these TFs were opening the chromatin around their bound sites.

**Figure 4.**
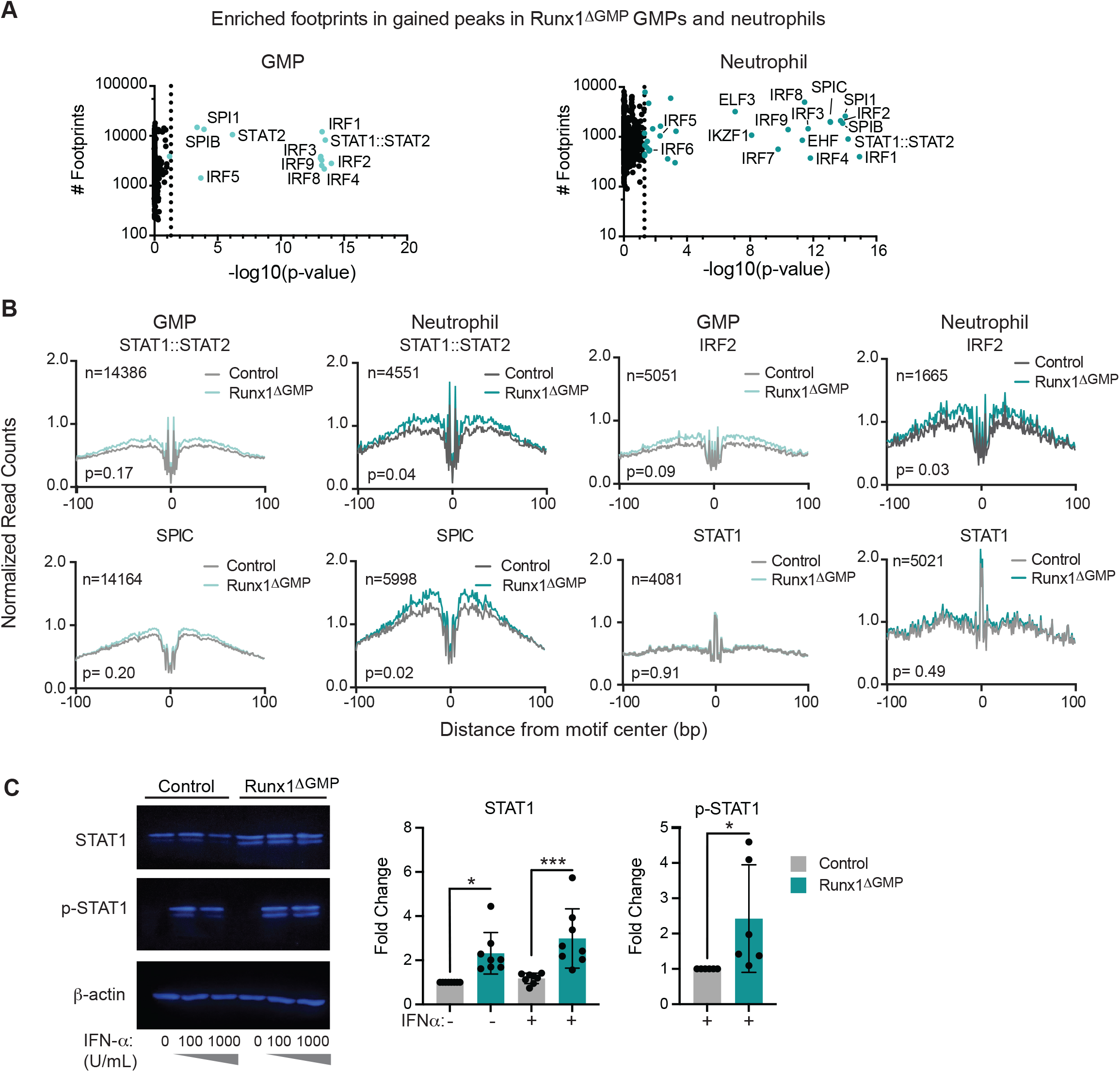
Chromatin opened following RUNX1 loss is enriched for footprints of TF effectors of type I IFN signaling. **A)** Scatter plots showing enriched digital TF footprints at regions of chromatin with increased accessibility in Runx1^ΔGMP^ neutrophils and GMPs relative to Controls. The number of footprints for each TF at regions of chromatin with increased accessibility is displayed on the y-axis for the Runx1^ΔGMP^cells. Colored circles indicate p<0.05; p-value calculated using biFET. **B)** Footprint profile plots for selected TFs showing average normalized read counts and p-values calculated by HINT-differential ^29^ using all peaks in Control and Runx1^ΔGMP^ cells. **C)** Western blot for STAT1 and phosphorylated (p) STAT1 plus β-actin control in Control and Runx1^ΔGMP^ neutrophils in the presence or absence of IFN-α. Summary of Western blots are on the right. Mean ± SD, dots indicate lanes quantified using ImageJ, ANOVA plus Tukey multiple comparison test for STAT1; two-tailed, unpaired t-test for p-STAT1. Representing 8 experiments, a total of 16 mice were analyzed. *P≤0.05, ***P≤0.001.

IRFs and STAT1::STAT2 are downstream effectors of type I IFN signaling, thus enrichment of their footprints suggests that type I IFN signaling was elevated in Runx1^ΔGMP^ neutrophils and GMPs. In contrast, the accessibility of chromatin around STAT1 footprints was not significantly increased in either Runx1^ΔGMP^ GMPs or neutrophils (Fig. 4B), therefore we infer that type II IFN signaling, which is mediated by the binding of STAT1 dimers to GAS sites, is not elevated to the same extent as type I IFN signaling. STAT3 and STAT5 footprints were also not enriched in gained peaks in Runx1^ΔGMP^ GMPs and neutrophils and the chromatin accessibility around their footprints was not increased (Fig. S5C), suggesting that cytokine signaling through the JAK/STAT pathway was not elevated. In conclusion, the predominant effect of RUNX1 loss in both GMPs and neutrophils was increased IRF and STAT1::STAT2 occupancy of sites that gained chromatin accessibility, suggesting that type I IFN signaling was elevated in both GMPs and neutrophils. These results are consistent with the RNA-seq data showing upregulation of genes associated with “cellular response to interferon-beta” and “response to interferon-beta” in Runx1^ΔGMP^ neutrophils and GMPs.

We considered why IRF and STAT1::STAT2 footprints were enriched in gained ATAC-seq peaks when RUNX1 was deleted. One possibility is that RUNX1 directly impedes the binding of IRFs and STAT1::STAT2 to adjacent sites. This does not, however, appear to be the case. Only 2.5% of the gained peaks in Runx1^ΔGMP^ neutrophils had RUNX1 footprints in Control neutrophils. Therefore, it is unlikely that RUNX1 directly interferes with the binding of IRF and STAT1::STAT2 at sites that gain accessibility in Runx1^ΔGMP^ cells. Another possibility is type I IFNs could be produced at higher levels by Runx1^ΔGMP^ cells. However, we found no evidence for increased production of IFN-α and/or IFN-β by neutrophils using CBA assays, or increased levels of IFN-α and/or IFN-β in the PB or BM with high sensitivity ELISA assays. Furthermore, expression from most of the *Ifna* and the *Ifnab* genes was undetectable in GMPs and neutrophils by RNA-seq. Therefore, increased type I IFN levels is unlikely to be the cause of increased IRF and STAT1::STAT2 occupancy. A third possibility is that RUNX1 represses the expression of signaling molecules in the type I IFN pathway, and RUNX1 loss sensitizes cells to respond to very low levels of IFNs. In support of this hypothesis, we found modestly increased STAT1 protein (but no increases in other STATs) in Runx1^ΔGMP^ neutrophils (Fig 4C, S6A-D), and RNA-seq read counts for type I IFN receptors (*Ifnar1, Ifnar2*), and two IRFs (*Irf1, Irf9)* were significantly higher in Runx1^ΔGMP^ GMPs (Fig. S6E.)

The failure to detect type I IFNs has been reported in other studies of cells with transcriptional and epigenetic type I IFN signatures ^31^. In some cases, it was shown that basal ISG expression is activated by non-canonical type I IFN signaling mediated by unphosphorylated (U) STATs (U-STAT1, U-STAT2) complexed with IFN response factor 9 (IRF9) ^32,33^. Unlike the canonical, IFN-stimulated pathway, tonic type I IFN signaling does not require activation of the type I IFN receptor and is unaffected by JAK inhibition ^31,33^. To determine whether the elevated basal expression of ISGs (*Stat1, Gbp2, and Irgm2)* in Runx1^ΔGMP^ neutrophils (Fig. 5A) was due to non-canonical IFN signaling we determined whether the JAK1/2 inhibitor ruxolitinib (Ruxo) could reduce the levels of mRNAs from the ISGs in Runx1^ΔGMP^ neutrophils in the absence of IFN-α stimulation. Ruxolitinib did not significantly decrease the basal levels of *Stat1, Gbp2*, or *Irgm2* mRNAs in Runx1^ΔGMP^ neutrophils in the absence of IFN-α, whereas the same concentration of ruxolitinib potently inhibited the increase in ISG mRNAs in response to IFN-α (Fig. 5B). A blocking antibody to IFNAR also failed to decrease the basal expression of ISGs in Runx1^ΔGMP^ neutrophils in the absence of IFN-α (Fig. 5C). We conclude that the increased expression of ISGs in Runx1^ΔGMP^ neutrophils is caused by elevated non-canonical type I IFN-induced signaling, possibly due to the increased expression of several proteins in the type I IFN signaling pathway, or increased chromatin accessibility of ISGs.

**Figure 5.**
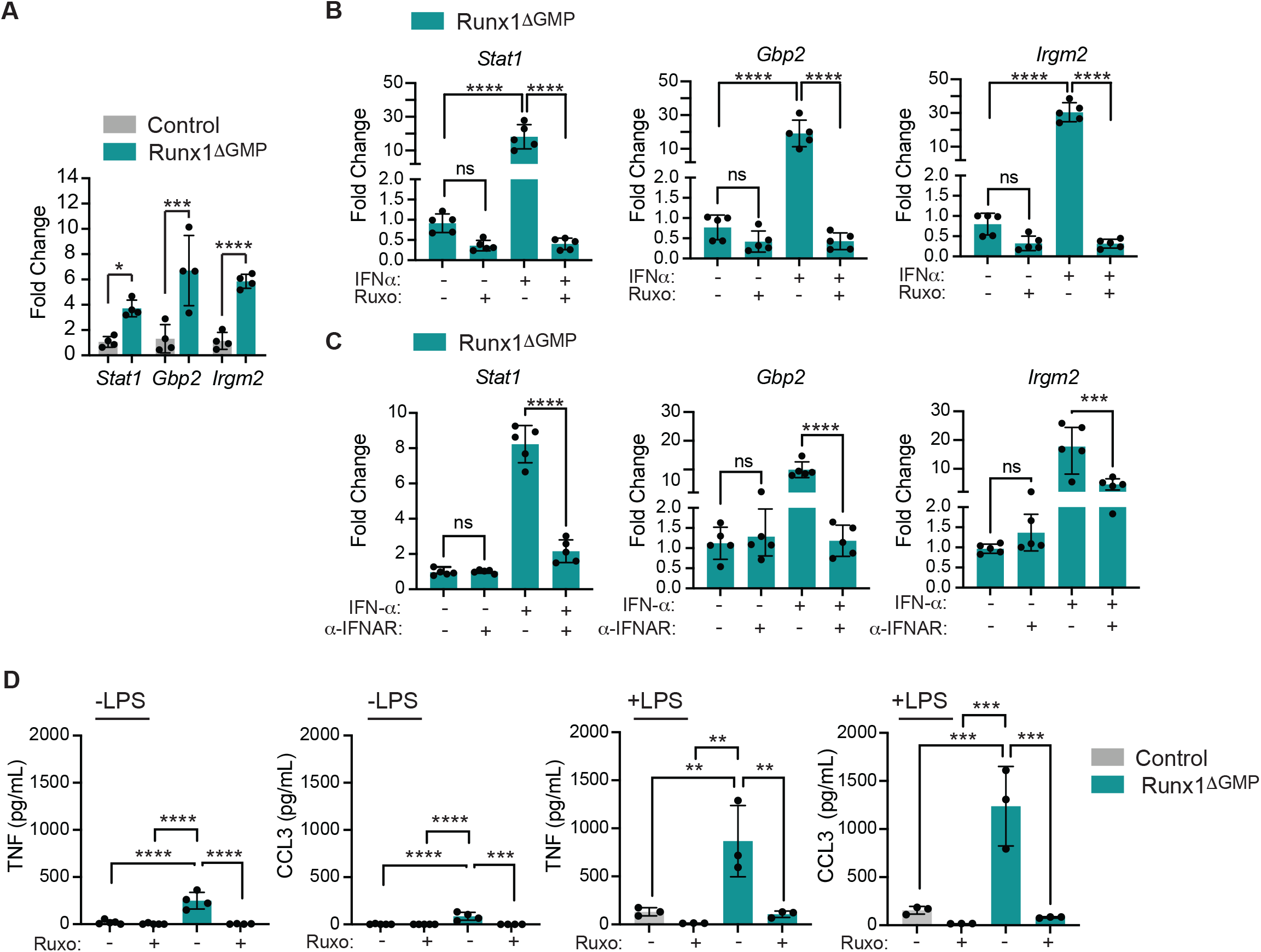
RUNX1 restrains tonic type I IFN signaling. **A)** RT-qPCR demonstrating baseline expression of three IGSs in Control and Runx1^ΔGMP^ neutrophils. Mean ± SD, two-tailed, unpaired t-test. Representing 2 experiments, total of 8 mice analyzed. **B)** The JAK inhibitor ruxolitinib (Ruxo, 20μM) blocks type I IFN signaling through the canonical pathway (+IFN-α), but not tonic signaling through a non-canonical pathway (-IFN-α) as measured by the expression of ISGs by RT-qPCR. All data are from Runx1^ΔGMP^ neutrophils. Representative of 2 experiments, total of 12 mice analyzed, mean ± SD, ANOVA plus Tukey multiple comparison test. **C)** Blocking antibody against the type I IFN receptor (α-IFNAR) does not decrease tonic signaling, as described in panel B for ruxolitinib. **D)** Ruxolitinib (20μM) decreases the production of TNF and CCL3 by neutrophils in response to LPS. Representative of 2 experiments, total of 15 mice analyzed, mean ± SD, ANOVA plus Tukey multiple comparison test. For all figures; *P≤0.05, **P≤0.01, ***P≤0.001, ****P≤0.0001, ns=not significant.

To determine if elevated type I IFN signaling was responsible for the hyper-response to LPS, we stimulated Runx1^ΔGMP^ neutrophils with LPS and tested whether ruxolitinib decreased the production of TNF and CCL3. TNF and CCL3 levels were significantly decreased, indicating that type I IFN signaling contributes to the overly exuberant response to LPS (Fig. 5D).

### RUNX1 restrains the chromatin accessibility and expression of retroelements

Elevated tonic type I IFN signaling has been associated with the expression of retroelements, including long terminal repeats/endogenous retroviruses (LTR/ERV) and long interspersed nuclear elements (LINE) ^34^. We examined ATAC-seq data from Runx1^ΔGMP^ GMPs and neutrophils to determine if the chromatin of retroelements was more accessible, and if so, whether motifs for IRFs and STATs were enriched in the opened chromatin. More ATAC-seq peaks in TEs were gained than lost in Runx1^ΔGMP^ GMPs and neutrophils, and there was an overall increase in the chromatin accessibility of TEs (Fig. 6A,B). The TEs with gained ATAC-seq peaks were LINE (LINE/L1) and LTR/ERV retroelements, whereas the short interspersed nuclear elements (SINE) and DNA transposons were significantly underrepresented in one or both cell types (Fig. 6C). Analysis of gained ATAC-seq peaks for enriched TF motifs identified several of the same TFs determined by genome-wide digital footprinting analyses, including motifs for multiple IRFs, STAT2, and SPI1 (Fig. 6D). The data suggest that increased tonic type I IFN signaling in Runx1^ΔGMP^ GMPs and neutrophils is associated with the increased chromatin accessibility of a subset of TEs, specifically the LINE and LTR/ERV retroelements.

**Figure 6.**
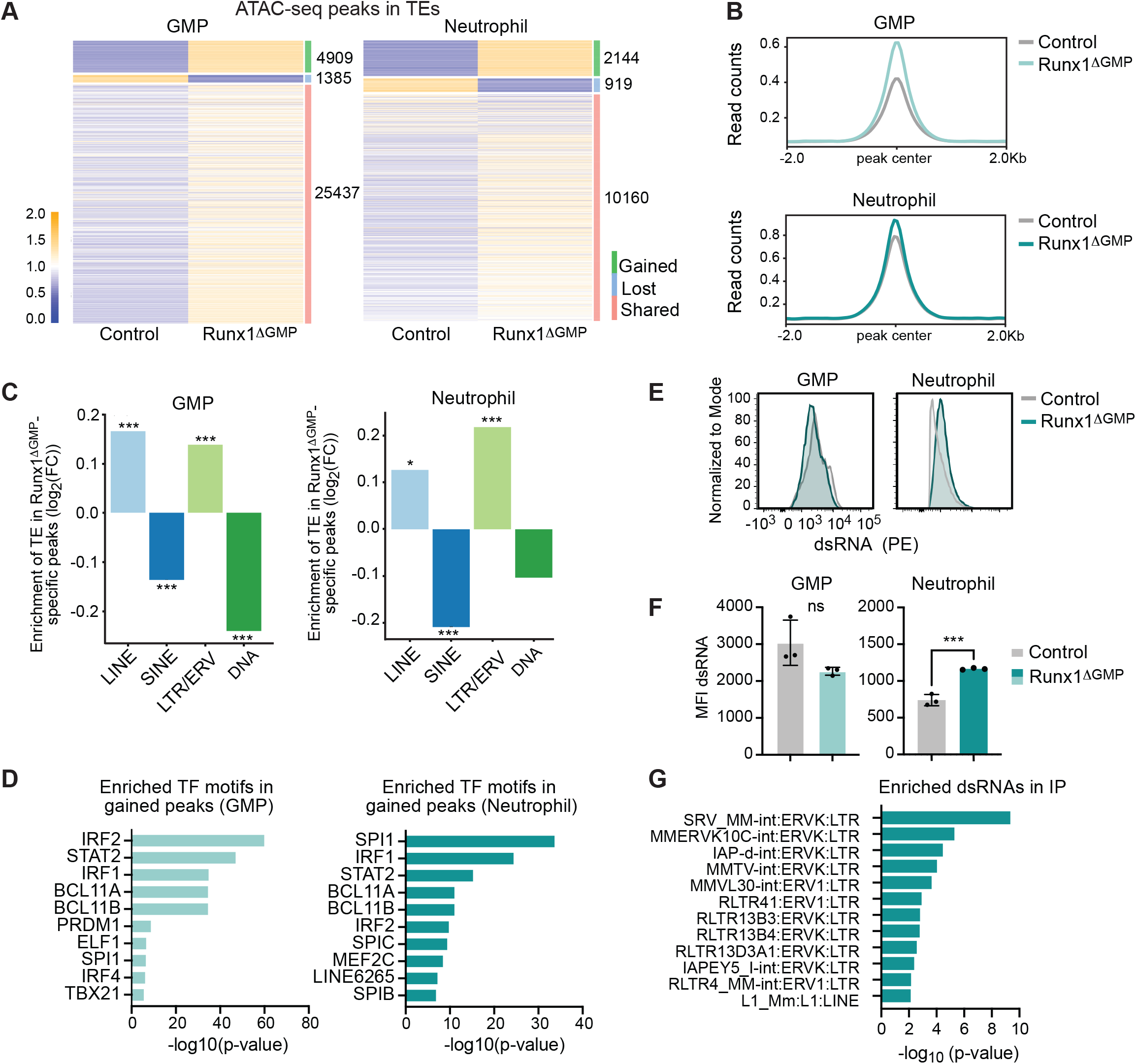
Loss of RUNX1 in GMPs derepresses TEs. **A)** Heatmaps of ATAC-seq peaks in TEs in GMPs and neutrophils. The number of gained, lost, and shared peaks are listed on the right side of each heat map. **B)** Averaged ATAC-seq peak profile plots (normalized to Bins Per Million mapped reads [BPM]) in TEs in Control and Runx1^ΔGMP^ neutrophils and GMPs.**C)** Enrichment or depletion of the different classes of TEs (LINE, SINE, LTR/ERV or DNA) in Runx1^ΔGMP^ neutrophils and GMPs. The y-axis represents the log2 fold change of the number of Runx1^ΔGMP^ specific peaks overlapping with TEs over the median number of the random selected peaks overlapping with TEs. Positive log2 fold change = enrichment, negative log2 fold change = depletion. **D)** Top 10 TF binding motifs enriched in Runx1^ΔGMP^ neutrophil- or Runx1^ΔGMP^ GMP-specific ATAC–seq peaks that overlapped with TEs. **E)** Representative histograms for the mean fluorescence intensities (MFI) of dsRNA in the dsRNA^+^ neutrophils or GMPs. **F)** Quantification of the relative MFI of dsRNA in the dsRNA^+^ neutrophils or GMPs. Statistics represent 2-tailed unpaired Student t tests. Representative of 2 experiments, total of 12 mice analyzed, ***P ≤ .001 **G)** IP of dsRNA with the 9D5 antibody followed by RNA-seq in Runx1^ΔGMP^ and Control neutrophils. Graph shows the dsRNA species detected and enriched in the IP from Runx1^ΔGMP^ neutrophils over Control neutrophils.

Retroelements are a source of dsRNA, and excess dsRNA is known to activate cytoplasmic dsRNA sensors that trigger the production of type I IFNs ^20^. To determine if increased chromatin accessibility in RUNX1 deficient cells resulted in the overproduction of dsRNA, we performed flow cytometry using a dsRNA antibody ^35^. The percentage of dsRNA^+^ cells, and the median fluorescence intensity (MFI) of dsRNA per cell were increased in Runx1^ΔGMP^ neutrophils although not in GMPs (Fig. 6E,F, S7). We immunoprecipitated and sequenced dsRNA from neutrophils with the same antibody used for flow cytometry. Immunoprecipitants of dsRNA from Runx1^ΔGMP^ neutrophils were enriched for multiple LTR/ERVK and one LINE subfamily (Fig. 6G), while other ERV subfamilies and dsRNA from DNA transposons were significantly depleted (Table S4). In conclusion, loss of RUNX1 elevates tonic type I IFN signaling, increases the chromatin accessibility of retroelements in GMPs and neutrophils, and increases the level of dsRNA produced from a subset of LTR/ERVKs and LINEs in neutrophils.

### Haploinsufficiency of RUNX1 may alter the properties of mouse and human neutrophils

FPDMM is caused by germline monoallelic *RUNX1* mutations ^36^. To determine if a monoallelic *Runx1* mutation is sufficient to alter inflammatory cytokine production, we analyzed neutrophils from mice heterozygous for a R188Q mutation, which affects a DNA-contacting residue (equivalent to R201Q in humans). The *Runx1*^*R188Q*^ mutation caused a small but significant increase in the amounts of TNF and CCL3 produced by neutrophils in response to LPS, demonstrating that monoallelic *Runx1* mutations are sufficient to cause a mildly hyperresponsive neutrophil phenotype (Fig. 7A). We then analyzed peripheral blood neutrophils isolated from two patients with FPDMM, and two unaffected family members by ATAC-seq. Neutrophils from each FPDMM patient and their family member were isolated and processed in parallel. FPD_21.1, a 9-year-old boy has a frameshift mutation (Tyr403Cysfs*153) in what is labeled as the transactivation domain by Clinical Genome Resource (ClinGen) ^37^ (Fig. 7B). Previously published analyses of RUNX1 activity in transactivation assays, however, showed that C-terminal truncation of murine RUNX1 at the human equivalent of Ser399 resulted in a protein was more transcriptionally active than full length RUNX1, hence this region has been functionally defined as a inhibitory domain ^38^. The Tyr403Cysfs*153 mutation is classified as likely pathogenic by ClinGen ^37^. Patient FPD_21.1 had a normal complete blood count (CBC) except for low platelets (73 K/mcL), elevated eosinophils (5.7%), and a small increase in immature granulocytes (0.6%) Patient FPD_21.1 had a persistent history of eczema. No known members of the FPD_21 family have had leukemia. FPD_52.3, an adult male has a pathogenic nonsense mutation (Arg201*) in the DNA-binding RUNT domain resulting in a non-functional protein (Fig. 7B). Patient FPD_52.3 also had a persistent history of eczema and environmental allergies. His CBCs were within normal ranges except for platelets (94 K/mcL) and eosinophils (14.1%). Three members of the FPD_52 family developed leukemia.

**Figure 7.**
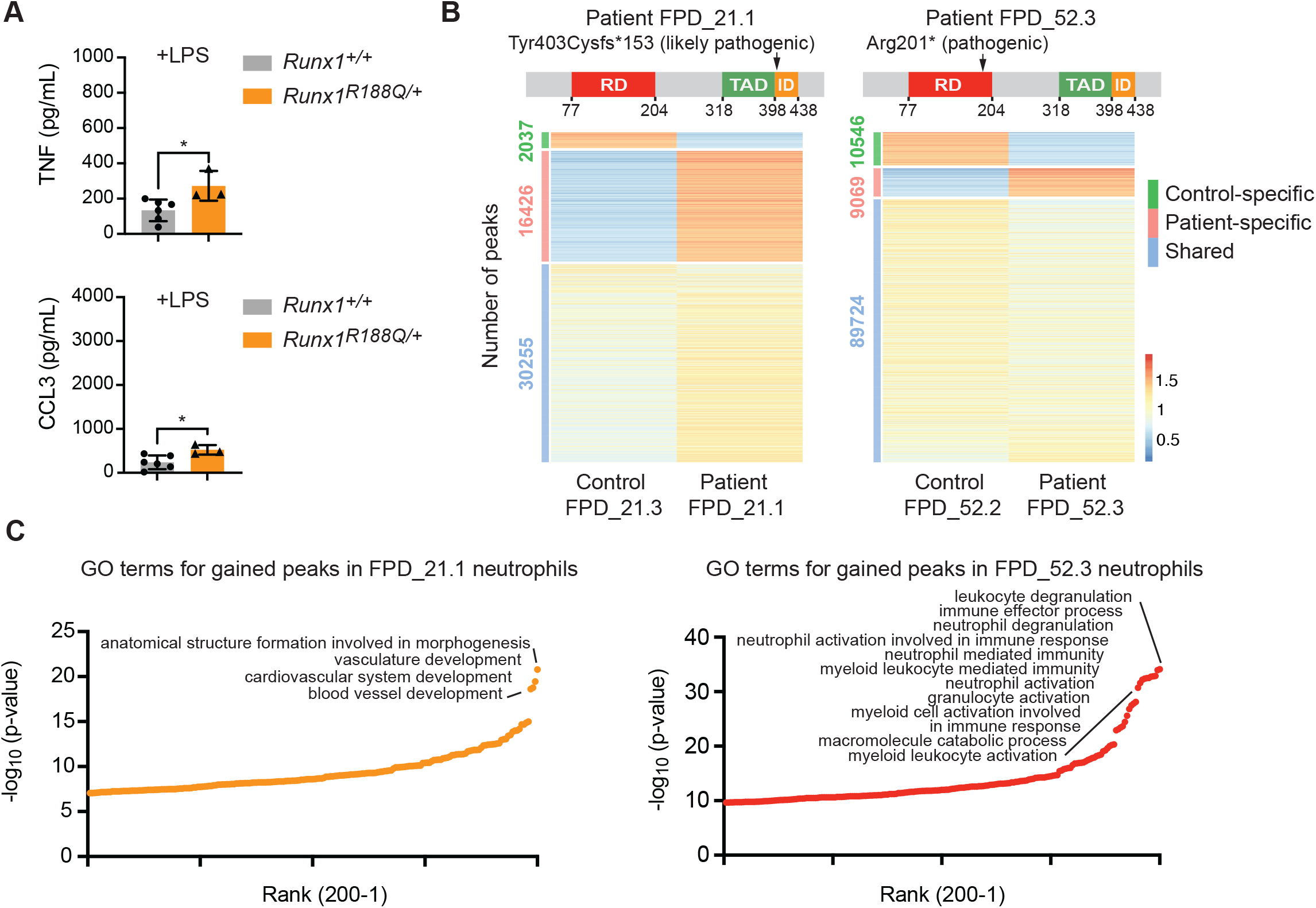
Haploinsufficiency of RUNX1 may alter the properties of mouse and human neutrophils. **A)** Absolute quantification by CBA of TNF and CCL3 in the supernatant of FACS-purified BM neutrophils from wild type or *Runx1*^*R188Q/+*^ mice stimulated with 100 ng/mL LPS. Mean ± SD, two-tailed, unpaired t-test, representative of 2 experiments, total of 9 mice analyzed, *P≤0.05. **B)** Top, schematic diagram showing location of the mutations in the RUNX1 protein in patients FPD_21.1 and FPD_52.3. ClinGen classifications of the mutations are indicated. RD, DNA-binding RUNT domain; TAD, transactivation domain, ID, inhibitory domain as defined by functional assays ^38,54^. Bottom, heatmaps of ATAC-seq signals for the two FPDMM patients and non-affected family members (Control). **C)** Enriched GO terms for patient-specific peaks. The top 200 GO terms are plotted.

The ratio of gained/lost peaks differed in neutrophils from FPD_21.1 and FPD_52.3 compared to their unaffected family members (Fig. 7B), as did the enriched GO Biological terms (Fig. 7C). The most highly enriched terms associated with peaks higher in FPD_21.1 than in his unaffected father (FPD_21.3) were related to vascular development, while those higher in FPD_52.3 relative to his unaffected wife (FPD_52.2) were related to neutrophil activation, suggesting FPD_52.3 neutrophils were in a more activated state (Fig. 7C). Terms related to type I IFN responses were not enriched in either patient or in control neutrophils. Many factors including age, environment, genetic backgrounds, health status, the nature of the *RUNX1* mutations, and the partial penetrance of inflammatory conditions in FPDMM patients could account for the difference between mice and patients, and between the two patients.

## DISCUSSION

We previously reported that pan-hematopoietic loss of RUNX1 resulted in neutrophils that hyper-respond to LPS, but that RUNX1 does not function in neutrophils *per se* to restrain the neutrophil response ^22^. Here we demonstrate that RUNX1 is required in a neutrophil precursor, either in the GMP or downstream of the GMP to restrain a pro-inflammatory epigenetic and transcriptional program in neutrophils. The pro-inflammatory program is characterized by a type I IFN signature, the de-repression of retroelements, and functional alterations in neutrophils, including the overproduction of inflammatory cytokines in response to TLR4 ligands.

The epigenetic rewiring in GMPs and neutrophils that results from RUNX1 loss has mechanistic parallels to trained immunity ^39^. In trained immunity, exposure to a pathogen induces epigenetic alterations in hematopoietic progenitors that are propagated to differentiated innate immune myeloid cells, causing them to respond more robustly to the same pathogen in a later encounter ^39^. This aspect of trained immunity provides obvious benefits to the infected host. However, trained immunity is also induced by endogenous molecules (danger-associated molecular patterns, or DAMPS) and continuous infections, and in these settings can be maladaptive, causing chronic inflammation. Mechanistic parallels between trained immunity and RUNX1 loss include: 1) epigenetic and/or transcriptional changes in GMPs, 2) acquisition of a type I IFN signature, and 3) neutrophils that hyper-respond to TLR ligands. For example, GMPs in mice fed a high fat Western diet acquired long-lasting transcriptional changes that were propagated to myeloid cells, causing them to hyper-respond to TLR ligands ^40^. In another mouse study, a brief period of experimentally induced periodontitis caused lasting maladaptive trained immunity, involving transcriptomic changes in HSPCs, an over-response of myeloid cells to the TLR4 ligand LPS, and systemic inflammation that increased the severity of arthritis, which is a frequent comorbidity in patients with periodontal disease ^41^. Treatment of mice with the fungal molecule β-glucan induced epigenetic and transcriptomic rewiring of GMPs and neutrophils that was associated with a type I IFN signature, and inhibition of type I IFN signaling abrogated the enhanced granulocyte responses ^42^. In humans, vaccination with tuberculosis vaccine bacillus Calmette-Guérin (BCG) caused long-term epigenetic changes that augmented neutrophils’ antimicrobial functions ^43^. We propose that loss or haploinsufficiency of RUNX1 causes a fixed state of maladaptive innate immunity. Further, we hypothesize that the intrinsic changes in HSPCs and neutrophils likely contribute to the increased incidence of inflammatory and allergic conditions in FPDMM patients, and possibly the elevated leukemia risk.

RUNX1 loss in GMPs results in the epigenetic and transcriptional de-repression of a subset of retroelements in neutrophils, specifically LINE and LTR/ERV. The mechanism by which this occurs is unclear. One possibility is that RUNX1 directly represses the chromatin accessibility of retroelements. However, since RUNX1 motifs are significantly depleted from sites that gain accessibility when RUNX1 is lost, repression of chromatin accessibility would more likely occur through interactions with other TFs at those sites. For example, BCL11A and BCL11B motifs are enriched in ATAC-seq peaks that are gained in retroelements when RUNX1 is deleted. RUNX1 is an important cofactor for BCL11B in both activating and repressing transcription during T cell development ^44^, therefore loss of RUNX1 could compromise a repressive activity of BCL11B on retroelements. The de-repression of retroelements could also be an indirect effect of RUNX1’s activity in restraining the expression of type I IFN signaling pathway molecules and tonic signaling ^13^. In this scenario, loss of RUNX1 would increase the occupancy and activity of IRFs and STATs on retroelements to increase their chromatin accessibility. The idea that increased activity of IRFs and STATs increases the chromatin accessibility and expression of retroelements is supported by our observation that the levels of dsRNA from ERVKs were elevated in RUNX1 deficient neutrophils, and ERVK is known to be regulated by IRFs and STATs ^45^. It also remains to be determined whether the increased expression of retroelements is only a consequence of elevated tonic type I IFN signaling and increased occupancy of IRFs and STATs, or if dsRNA produced from retroelements functions in a positive feedback loop to promote the production of type I IFNs that further activate type I IFN signaling in an autocrine or paracrine manner. We were unable to detect elevated levels of type I IFNs in the BM of Runx1^ΔGMP^ mice, with the caveat that their concentrations may be below the limit of detection. Therefore, we currently have no evidence to support this last mechanism.

Mutations in several genes associated with clonal hematopoiesis of indeterminate potential (CHIP) increase the inflammatory properties of myeloid lineage cells, specifically macrophages and neutrophils ^46^. CHIP is caused by somatic mutations acquired by HSCs that confer a selective advantage to the mutated HSC, causing it to preferentially expand in the BM relative to unmutated HSCs ^47,48^. CHIP elevates the risk of cardiometabolic diseases by increasing the inflammatory properties of myeloid lineage cells that differentiate from the mutated HSCs ^46^. To our knowledge, the de-repression of TEs in myeloid lineage cells with CHIP mutations has not been demonstrated, but we believe is worth examining, particularly as the most common CHIP mutations involve epigenetic regulators (DNMT3A, TET2, ASXL1) and a kinase in the type II IFN pathway (JAK2) ^46^.

Our studies focused on neutrophils, but the regulation of type I IFN signaling and chromatin accessibility of retroelements by RUNX1 is not necessarily confined to this lineage. Previous work demonstrated that loss of RUNX1 function in lung alveolar epithelial cells, B cells, and BM cells increased the production of type I IFNs and/or ISGs ^12-14^. Therefore, FPDMM patients may have a more generalized dysregulation of retroelement chromatin accessibility and inflammation involving multiple tissues in which RUNX1 is expressed that could contribute to their inflammatory conditions.

## METHODS

### Mice

Runx1^ΔHSC^, Runx1^ΔLYM^, or Runx1^ΔGMP^ mice were created by breeding *Runx1*^*f/f*^ (*Runx1*^*tm1Spe*^) mice ^49^ with Vav1-Cre (Tg(Vav1-cre)1Graf) ^50^, Rag1-Cre (Rag1^tm1(cre)Thr^) ^51^, or Cebpa-Cre (Cebpa^tm1.1(cre)Touw^) ^52^ mice. *Rag2*^*-/-*^ mice (B6.Cg-*Rag2*^*tm1*.*1Cgn*^/J) were purchased from JAX. *Cd14*^*-/-*^ mice (B6.129S4-Cd14^tm1Frm/J^) ^27^ were obtained from Carla R. Scanzello. *Runx1*^*R188Q/+*^ *(C57BL/6J-Runx1*^*<tm1Lhc>R188Q*^) knock in mice will be described elsewhere. Male and female mice ages 6 to 12 weeks were used in all experiments. Mice were handled according to protocols approved by the University of Pennsylvania’s Institutional Animal Care and Use Committee and housed in a specific pathogen-free facility.

### Flow cytometry and cell sorting

A full list of antibodies is provided in Table S1. Flow cytometry was performed on an LSR II, and data analyzed using FlowJo software. The lineage panel in Fig. S7 includes CD3, CD11b, B220, Gr-1, Nk1.1, and Ter119. Cells analyzed by intracellular flow were fixed and permeabilized using Cytofix/Cytoperm (Benton Dickinson (BD)) prior to intracellular staining in the perm/wash buffer (BD). For dsRNA intracellular flow, cells were permeabilized with 3% PFA for 15 min on ice, washed two times with FACS buffer (2% FBS in 1x PBS) and then permeabilized with 0.1% Saponin for 15 min on ice. The cells were then incubated with 9D5 dsRNA rabbit IgG mAb (1:500) in 0.1% Saponin for 15 min on ice, washed twice with 0.1% Saponin, and then incubated with Donkey anti-Rabbit IgG (minimal x-reactivity) PE antibody (1:400) for 15 min on ice. The cells were rinsed three times with FACS buffer. Gating schemes for intracellular flow assays are provided in Figs. S1A and S7. A BD fluorescence-activated cell sorter (FACS) Aria II was used to sort cells at 482.63 kPa (70 psi) using a 70-μm nozzle. Gating schemes for purified neutrophils and GMPs are provided (Fig. S7).

### Statistical analysis

Unless otherwise indicated, all statistical analyses were performed using Prism/GraphPad.

### FPDMM patient data

FPDMM patient data was obtained following informed consent under the clinical study entitled “Longitudinal Studies of Patients with FPDMM” (ClinicalTrials.gov identifier: NCT03854318).

### Data and code availability

The data discussed in this publication have been deposited in NCBI’s Gene Expression Omnibus ^53^ and are accessible through GEO Series accession number GSE221427 (https://www.ncbi.nlm.nih.gov/geo/query/acc.cgi?acc=GSE221427).

## COMPETING INTEREST STATEMENT

The authors declare no competing interests.

## ACKNOWLEDGMENTS

We thank William Murphy, Jennifer Jakubowski, and Shifu Tian in the Flow Cytometry and Cell Sorting Resource Laboratory for cell sorting assistance. This work was supported by R01HL091724 (NAS), U01HL100405 (NAS), 1F30DK128926-01A1 (A.U.Z.), T32 HL7439-43 (A.U.Z.), 1F31HL150952-01 (EDH), T32 HD083185 (EDH), the RUNX1 Research Program (NAS), and the Intramural Research Program, National Human Genome Research Institute, NIH (EB, JD, PPL).

## AUTHOR CONTRIBUTIONS

N.A.S., A.U.Z., and W.Y. conceived the project and wrote the manuscript. All authors discussed the results and revised the manuscript. A.U.Z., D.Y, E.D.H., D.Y., J.R. and J.D. performed the experiments. W.Y. and A.U.Z. generated the genomics data. W.Y., A.U.Z., and E.D.H. performed bioinformatics analyses. M.H.A, L.H.C. and I.P.T. provided mice. A.J.M, W.T., K.T. provided advice. P.P.L. and J.D. provided FPDMM patient samples.

## Notes

### Competing Interest Statement

The authors have declared no competing interest.

