## Supplemental Table 1 for "RUNX1 is required in granulocyte-monocyte progenitors to attenuate inflammatory cytokine production by neutrophils"

| **Antibody** | **Clone** | **Fluorophore** | **Supplier** | **RRID #** | **Dilution** |
| --- | --- | --- | --- | --- | --- |
| c-Kit | 2B8 | FITC | Biolegend | AB_313215 | 1:200 |
| CD3e | 145-2C11 | APC | Biolegend | AB_312677 | 1:200 |
| CD3e | 145-2C11 | PE | Biolegend | AB_312672 | 1:200 |
| CD4 | RM4-5 | BV 421 | Biolegend | AB_2563052 | 1:200 |
| CD8 | 53-6.7 | PE-Cy7 | Biolegend | AB_312761 | 1:200 |
| CD14 | Sa14-2 | PE | Biolegend | AB_940584 | 1:200 |
| CD14 | Sa14-2 | Unconjugated | Biolegend | AB_940588 | 1:200 |
| CD14 | 4C1/CD14 | Unconjugated | BD Biosciences | AB_396926 |  |
| CD16/32 | 93 | APC-Cy7 | Biolegend | AB_2104158 | 1:200 |
| CD19 | 1D3 | APC | Thermo Fisher | AB_1659676 | 1:200 |
| CD19 | 1D3 | PE | Biolegend | AB_2629817 | 1:200 |
| CD45.1 | A20 | APC-Cy7 | Biolegend | AB_313505 | 1:200 |
| CD45.1 | A20 | PE-Cy7 | Thermo Fisher | AB_469629 | 1:200 |
| CD45.2 | 104 | FITC | Thermo Fisher | AB_465062 | 1:200 |
| B220 | RA3-6B2 | APC | Biolegend | AB_312997 | 1:200 |
| CD150 | TC15-12F12.2 | PE-Cy7 | Biolegend | AB_439797 | 1:200 |
| F4/80 | BM8 | FITC | Biolegend | AB_893500 | 1:200 |
| Gr-1 | RB6-8C5 | APC | Biolegend | AB_313377 | 1:200 |
| Gr-1 | RB6-8C5 | PerCP-Cy5.5 | Thermo Fisher | AB_906247 | 1:200 |
| IFNAR-1 | MAR1-5A3 | Unconjugated (blocking Ab) | Bio X Cell | AB_2687723 |  |
| Ly-6G | 1A8 | APC | Biolegend | AB_2227348 | 1:200 |
| Ly-6G | 1A8 | PE- Cy7 | BD Biosciences | AB_1727562 | 1:200 |
| Ly-6G | 1A8 | APC-Cy7 | BD Biosciences | AB_1727561 | 1:200 |
| Mac-1 | M1/70 | APC | Biolegend | AB_312795 | 1:200 |
| Mac-1 | M1/70 | APC-Cy7 | BD Biosciences | AB_396772 | 1:200 |
| Nk1.1 | PK136 | APC | Biolegend | AB_313397 | 1:200 |
| Rat IgG2a, κ Isotype Ctrl | RTK2758 | PE | Biolegend | AB_326530 | 1:200 |
| Mouse IgG1, κ Isotype Ctrl | monoclonal antibody | Unconjugated | BD Biosciences | AB_10050442 |  |
| Sca-1 | D7 | PerCP-Cy5.5 | Thermo Fisher | AB_914372 | 1:200 |
| Sca-1 | E13-161.7 | PE | Biolegend | AB_756193 | 1:200 |
| Siglec F | E50-2440 | PE | BD Biosciences | AB_394341 | 1:200 |
| Siglec F | E50-2440 | APC-Cy7 | BD Biosciences | AB_2732831 | 1:200 |
| Streptavidin |  | PerCP-Cy5.5 | Biolegend | AB_2716577 | 1:200 |
| Ter119 | TER-119 | APC | Biolegend | AB_313713 | 1:200 |
| TNF-alpha | MP6-XT22 | Pacific Blue | Biolegend | AB_893639 | 1:100 |
| dsRNA | 9D5 | Unconjugated | Absolute Antibody | Cat. No: Ab00458-23.0 | 1:500 |
| Donkey anti- rabbit IgG | Poly4064 | PE | Biolegend | AB_2563484 | 1:400 |
| Anti-Histone H3 (acetyl K27) antibody | polyclonal antibody |  | Abcam | AB_2118291  Cat. No: ab4729 |  |
| β-Actin | C4 |  | Santa Cruz Biotechnology | AB_2714189  Cat. No: sc-47778 HRP | 1:1000 |
| GAPDH | D4C6R |  | Cell Signaling Technology | AB_2756824  Cat. No: 97166 | 1:1000 |
| STAT1 |  |  | Cell Signaling Technology | AB_2198300  Cat. No: 9172 | 1:1000 |
| pSTAT1 (Tyr701) | 58D6 |  | Cell Signaling Technology | AB_561284  Cat. No: 9167 | 1:1000 |
| STAT2 | D9J7L |  | Cell Signaling Technology | AB_2799824  Cat. No: 72604 | 1:1000 |
| pSTAT2 (Tyr689) |  |  | Millipore | AB_2198439  Cat. No: 07-224 | 1:1000 |
| STAT3 |  |  | Cell Signaling Technology | AB_2629499  Cat. No: 12640 | 1:1000 |
| pSTAT3 (Tyr705) | D3A7 |  | Cell Signaling Technology | AB_2491009  Cat. No: 9145 | 1:1000 |
| STAT5 | D2O6Y |  | Cell Signaling Technology | AB_2737403  Cat. No: 94205 | 1:1000 |
| pSTAT5 (Y694) |  |  | Cell Signaling Technology | AB_2315225  Cat. No: 9351 | 1:1000 |
| IRF9 | D9I5H |  | Cell Signaling Technology | AB_2798964  Cat. No: 28845 | 1:1000 |
| DAPI |  |  | Thermo Fisher | Cat. No: D1306 |  |
| LIVE/DEAD Fixable Aqua |  |  | Thermo Fisher | Cat. No: L34957 |  |

**Supplementary Table 1.** List of all antibodies used for flow cytometry, western blot analysis, and H3K27ac ChIP-seq. For each antibody, the clone, fluorophore, dilution, manufacturer, and antibody registry number (or manufacturer’s catalog number when no RRID number is available) are provided.
