## Supplemental Methods and Figures for "RUNX1 is required in granulocyte-monocyte progenitors to attenuate inflammatory cytokine production by neutrophils"

**SUPPLEMENTARY METHODS:**

**Bone Marrow Harvest:** Bone marrow (BM) cells were harvested by spinning isolated bone at 12,000 x g for 1 min, filtering through a 70-µm filter, and resuspending in cold PBS with 2% heat-inactivated FBS. Red blood cells were lysed in ACK Lysing Buffer (Gibco) for 2 min on ice.

**Transplant analyses:** B6.SJL-*Ptprc*$^{a}$*Pepc*$^{b}$/BoyJ (CD45.1) mice were subjected to a split dose of 1000 cGy, 3 h apart. Each recipient received a 10:1 ratio of BM cells from control (CD45.1/2) and Runx1$^{\Delta HSC}$ mice (CD45.2) by retro-orbital injection. We assessed donor (CD45.2) and competitor (CD45.1/2) engraftment in peripheral blood at weeks 4, 16, and 20. Mice were sacrificed at 24 weeks post-transplant and bone marrow was harvested for final engraftment and cytometric bead array analyses.

***Ex vivo* culture and stimulation:** Following isolation of neutrophils via FACS, cells were rested for two hours at 37°C in Hank's media consisting of 1x Hank's Balanced Salt Solution (Gibco) with 25 mM HEPES and 10% heat inactivated fetal bovine serum (FBS; Gemini Bio-Products) unless otherwise indicated. Cells were stimulated at 37°C with LPS (10 ng/mL or 100 ng/mL) (E. coli O111:B4, Imgen Technologies) in Hank's media. For the CD14 blocking experiments, neutrophils were incubated prior to LPS stimulation for 30 min at 37°C with 5, 15 or 50 µg/mL of function-blocking anti-CD14 or isotype control. For the ruxolitinib experiments, when indicated, neutrophils were incubated for 1 hour with 20 µM of ruxolitinib (Selleck Chemicals) prior to LPS stimulation. For intracellular flow assays, stimulation media also included Brefeldin A (BD GolgiPlug).

**Cytokine quantification:** 200,000 neutrophils were plated in a total volume of 100 µL into 96 well plates. After 8 hours of LPS stimulation, cells were pelleted, and supernatants frozen until analyzed. Absolute multiplex quantification of an 8-factor panel of cytokines, chemokines, and growth factors was performed using the Cytometric Bead Array (CBA) mouse soluble protein flex sets (BD) according to the manufacturer's instructions. A sigmoidal 4-parameter logistic regression was used to fit a standard curve and interpolate unknown concentrations.

**Real-time quantitative reverse transcriptase-PCR (RT-qPCR)**: Isolated neutrophils (EasySep™ Mouse Neutrophil Enrichment Kit) were incubated in the presence of 20 μM of ruxolitinib (Selleck Chemicals), 1 μ/mL anti-mouse IFNAR-1 mAB, or vehicle, or isotype control for 30 min- 1 hr. Cells were then stimulated with 1,000 U/mL INF-α or vehicle for 1-2 hrs. Total RNA was isolated from neutrophils (RNeasy Mini Kit, Qiagen) and quantified by Nanodrop. Total RNA was reverse transcribed into cDNA (ProtoScript® II First Strand cDNA Synthesis Kit, NEB). Relative STAT1, CD14, IRGM2, and GBP2 mRNA levels were determined using Power SYBR Green Master Mix (Thermo Fisher Scientific). Cyclophilin was used as an endogenous control. qPCR primers are: CD14 F: GGCGCTCCGAGTTGTGACT; CD14 R: TACCTGCTTCAGCCCAGTGA; STAT1 F: GCCTCTCATTGTCACCGAAGAAC; STAT1 R: TGGCTGACGTTGGAGATCACCA; Cyclophilin F: ATGGCAAATGCTGGACCAA; Cyclophilin R: GCCATCCAGCCATTCAGTCT; IRGM2 F: CCCCTTCTTTCACGGCAGT; IRGM2 R: GGCAGTTGAGTCACCTGAGG; GBP2 F: CTGCACTATGTGACGGAGCTA; GBP2 R: CGGAATCGTCTACCCCACTC.

**Western Blot**: Neutrophils were starved in 0.05% BSA for 2-4 hrs, and then stimulated with vehicle or IFN-α for indicated time points and snap-frozen in dry ice. Cell pellets were lysed in LDS loading buffer and sonicated for homogenization. Samples were resolved by SDS-PAGE and transferred to a nitrocellulose membrane. For all primary phospho-antibody blots, membranes were blocked with 5% BSA (BP1600-100, Fisher Bioreagents) in TBS-T, while other primary antibody blots were blocked with 5% non-fat milk (sc2325, Santa Cruz). Membranes were incubated with primary antibodies overnight in a cold room (complete list of antibodies listed in Table S1). Following primary antibody blots, membranes were washed with TBS-T, and then incubated with HRP-conjugated secondary antibody for 1 hr at room temperature. After washing, membranes were developed with ECL (#34095, Thermo Scientific). In certain cases, Western blots were stripped and reprobed with a second set of primary antibodies. Immunoblots were processed and developed by KwikQuant imager (Kindle Biosciences, LLC). Quantification of western blots was performed using ImageJ software.

**H3K27ac ChIP-Seq:** FACS purified neutrophils ($CD11b^{+}SiglecF^{-}F4/80^{-}Ly6G^{+}$) were fixed and crosslinked by 1% formaldehyde in 1× Fixing Buffer for 5 min at room temperature according to the vendor's protocol (Covaris). Cells were then stored at -80℃. Crosslinked cells were thawed on ice (100K cells) and resuspended in 1×Shearing Buffer and sonicated with Covaris E220 for 720s using the following settings: 5% duty factor, 105W Peak Incident Power, and 200 cycles per burst. 10% of sheared chromatin was used as the input and the remaining chromatin was divided into two equal aliquots for immunoprecipitation (IP) (50,000 cells per IP). IPs were performed using ChIP-IT high sensitivity kit (Active Motif). IP and input samples were treated with RNase A followed by proteinase K. Crosslinking was reversed by incubation overnight at 65 ℃ and DNA was purified using a MinElute PCR purification kit (Qiagen). All IP DNA and 1-2 ng of input DNA were used for library preparation with the ThruPLEX DNA-Seq kit and Smarter DNA single index kit (Takara). 13 and 9 cycles were used for IP DNA and input DNA, respectively at step 5. Following library amplification, the libraries were bead purified. The concentrations were measured using both Qubit and KAPA qPCR. Agilent Bioanalyzer 2100 was used to check the quality of libraries. Libraries were sequenced on Illumina HiSeq 2500 sequencer in single-end mode with read length of 75bp.

**H3K27ac ChIP-Seq data processing:** Sequencing reads were demultiplexed using Bcl2Fastq v2.20 then trimmed and filtered for quality using Trim Galore (Martin 2011) with the following settings: fastqc, and trim1. Reads were then aligned to the mouse genome (mm10) using bowtie 2.3.5.1 [55,56]. Only uniquely mapped reads with fewer than 2 mismatches were used for downstream analyses. Samtools v.1.1 [57] was used to convert SAM files to BAM files, and Sambamba v0.6.6 [58] was used to filter out duplicates, multi-mappers, reads mapped to ChrM or blacklist regions, and unmapped reads. MACS2 2.1.4 [59] was used for peak calling with the following parameters: narrow, q: 0.05. Control or $Runx1^{\Delta GMP}$ specific differential peaks were called if the RPKM fold change between Control and Runx1$^{\Delta GMP}$ peaks was greater than 2. Merged replicates were used to create bigwig files for visualization using Deeptools v3.3.0 [60]). The following parameters were used: normalized to reads per genomic content (RPGC), effective genome size: 2,308,125,349 bp, ignore for normalization: ChrX, min fragment length: 20, bin size: 10. IGV or UCSC genome browser was used for visualization. Deeptools was also used to plot regions of differential peaks. GREAT 4.0.4 [61,62] was used for linking peak regions to genes and subsequent gene ontology annotation using the following parameters: species assembly: mm10; association rule: basal+extension: 5000 bp upstream, 1000 bp downstream, 1,000,000 bp max extension, curated regulatory domains included.

**Bulk ATAC-seq:** Neutrophils from human patient/control peripheral blood were isolated using the EasyStep Human Neutrophil Isolation Kit (Stem Cell). Patient and control neutrophils were isolated and processed side-by-side on the same day. 50,000 neutrophils were collected and washed with cold PBS. Cell pellets were resuspended in 50 µl of cold lysis buffer (10 mM Tris-HCl, pH 7.4, 10 mM NaCl, 3 mM $MgCl_2$ and 0.1% Tween-20, 0.1% NP-40, 0.01% Digitonin, 1% BSA) and incubated on ice for 4 min. Lysis was halted with the addition of 50 µL of wash buffer (10 mM Tris-HCl, pH 7.4, 10 mM NaCl, 3 mM $MgCl_2$ and 0.1% Tween-20, 0.1%, 1% BSA). Samples were centrifuged at 500 x g, 4 °C for 5 min. Nuclei pellets were resuspended in 50 µl of wash buffer and immediately centrifuged at 500 x g, 4 °C for 5 min. Nuclei pellets were resuspended in 50 µl of transposition reaction mix (1× Tagment DNA Buffer, 2.5 µl of Tagment DNA Enzyme 1) and incubated for 30 min at 37 °C. Subsequent steps of the protocol were performed as previously described [63]. Libraries were purified using a Qiagen MinElute Gel Purification kit for mouse samples and SPRI-Select bead purification for human samples. The concentrations were measured using both Qubit and KAPA qPCR. Agilent Bioanalyzer 2100 was used to determine the quality of libraries. Libraries were sequenced on the Illumina HiSeq 2500, with 75-bp paired-end reads. Each sample had two biological replicates.

**Bulk ATAC-seq data processing for mouse samples:** Sequencing reads were demultiplexed using Bcl2Fastq v2.20 then trimmed and filtered for quality using Trim Galore v0.6.4 [64] with the following settings: fastqc, paired, and trim1. Reads were then aligned to the mouse genome (mm10) using bowtie 2.3.5.1 [55,56]. Only uniquely mapped reads with fewer than 2 mismatches were used for downstream analyses. Samtools v.1.1 [57] was used to convert SAM files to BAM files, and Sambamba v0.6.6 [58] to filter out duplicates, multi-mappers, reads mapped to ChrM or blacklist regions, and unmapped reads. MACS2 2.2.7.1 [59] was used for peak calling using the following parameters: BAMPE, q: 0.05. Peaks from Control and Runx1$^{\Delta GMP}$ were merged if there is at least 1bp overlap between two peaks. Reads Per Kilobase Million (RPKM) of a peak was then calculated using bedtools v2.25 [65]. Peaks with RPKM less than 0.5 in both Control and Runx1$^{\Delta GMP}$ cells were filtered from downstream analysis. Control or Runx1$^{\Delta GMP}$ specific peaks were called if the RPKM fold change between Control and Runx1$^{\Delta GMP}$ peaks was greater than 2. Merged replicates were used to create bigwig files for visualization using Deeptools v3.3.0 [60] and the following parameters: normalized to Bins Per Million mapped reads (BPM), bin size: 50. IGV or UCSC genome browser was used for visualization. Deeptools was also used to plot regions of differential peaks. GREAT 4.0.4 [61,62] was used for linking peak regions to genes and subsequent gene ontology annotation using the following parameters: species assembly: mm10; association rule: basal+extension: 5000 bp upstream, 1000 bp downstream, 1,000,000 bp max extension, curated regulatory domains included. Homer v4.11 [66] was used for genomic annotation of peak regions.

**Bulk ATAC-seq data processing for human samples:** Human data were processed as described with the following changes. Reads were aligned to the human genome (hg38) using Bowtie 2.3.5.1 [55,56]. MACS2 2.2.7.1 [59] was used for peak calling using the following parameters: BAMPE, q: 0.05. Peaks from the patient and control samples were called individually. For downstream analysis, we compared the patient and unaffected family members individually. The peaks from the patient sample and the corresponding control sample were merged for each comparison and RPKM per merged peak was calculated using BEDtools. Peaks with RPKM less than 0.5 in both control and patient were filtered from downstream analysis. Control or patient specific peaks were called if the RPKM fold change between control and patient peaks was greater than 2. Individual samples were used to create bigwig files for visualization. GREAT 4.0.4 [61,62] was used for linking peak regions to genes and subsequent gene ontology annotation using the following parameters: species assembly: hg38; association rule: basal+extension: 5000 bp upstream, 1000 bp downstream, 1,000,000 bp max extension, curated regulatory domains included.

**Footprinting analysis and motif analysis:** ATAC-seq footprinting analysis was performed using the Regulatory Genomics Toolbox (RGT) and HMM-based IdeNtification of Transcription factor footprints (HINT) software [29]. In brief, footprints were called from regions of chromatin accessibility (peaks on merged replicates) for each sample. Called footprints were then matched to TF motifs using the JASPAR 2020 vertebrate motif database. Differential activity of transcription factors, as well as plots of each transcription factor footprint, were determined using HINT-differential. p-value and read counts were calculated using RGT HINT-differential for each transcription factor. Enriched transcription factor footprints at regions of chromatin with increased accessibility and corresponding p-values were determined using BiFET v.1.16.0 software [30]. TF footprints were extracted for more in depth analysis. TF motif scores were determined by the RGT toolbox [67]. ATAC-seq motif analysis was done as follows: TF motifs were first scanned on all peaks by R package motifmatchr and a motif hit was called on a peak if the p-value was less than $10^{-5}$. The TF motif was enriched on a list of Runx1$^{\Delta GMP}$ or Control specific peaks if the p-value of a binomial test, comparing the TF motif on the rest of the peaks, was less than 0.05.

**Enrichment of transposable elements:** An ATAC-seq peak is associated with a TE (annotated by RepeatMasker, Reference: Smit, AFA, Hubley, R & Green, P. *RepeatMasker Open-4.0*. 2013-2015 http://www.repeatmasker.org) if there is at least 1 base pair overlap between them. Enrichment of TEs were performed by the following permutation test for each TE class and family, respectively: For a given TE class (or family), we counted the number of gained peaks in Runx1$^{\Delta GMP}$ cells that overlapped with any TE belonging to the given TE class (or family). We randomly selected the same number of peaks from the rest of the peaks (lost peaks in Runx1$^{\Delta GMP}$ cells or stable peaks) and counted the number of peaks that overlapped with the given TE class (or family). This process was repeated 1000 times to generate a null distribution of the number of peaks overlapped with TE. The empirical p-value was then calculated as the number of times the permutation yielded values greater than the number of gained peaks overlapped with the TE class (or family), divided by 1000.

**Cytosolic dsRNA immunoprecipitation and sequencing**: dsRNA was immunoprecipitated as previously described [68]. Briefly, Protein G Dynabeads and Protein A Dynabeads (1:1) were washed and resuspended in NET-2 buffer. 5 µg of 9D5 dsRNA rabbit IgG mAb was bound to the beads for 1-2 hours in the cold room on a shaker. Six million FACS purified neutrophils were lysed for 10 min by end-over-end rotation in the cold room in cytosolic lysis buffer. The cell lysate was centrifuged at 980 x g for 3 min at 4℃. The centrifugation steps were repeated for a total of three times. The cytosolic supernatant was then transferred to a fresh tube and spun at 17,000 x g for 10 min. For immunoprecipitation, lysate was diluted 1:4 with NET-2-TurboDNase buffer. 95D-Dynabeads were added to the lysate and end-to-end rotated for 2 hr at 4 °C. Following magnetic separation, beads were washed with high salt washing buffer and then NET-2 buffer. 95D-bound dsRNA was extracted with Trizol reagent and purified using Direct-zol™ RNA Miniprep Kit (Zymo Research). RNA samples were ribo-depleted with Ribominus™ Eukaryote v2 kit (Thermo Fisher Scientific). Libraries were prepared using the NEBNext Ultra II Directional RNA Library Prep Kit for Illumina. Libraries were sequenced on Illumina HiSeq 2500 sequencer in paired-end mode with read length of 75bp.

**dsRNA-seq data processing:** Raw dsRNA-seq sequencing data (.bcl files) was converted into Fastq files and de-multiplexed using Bcl2Fastq v2.20 software. The data in fastq file format was processed with the toolkit SQuIRE [69]. The raw reads were aligned to the mm10 reference genome and TEs were annotated using RepeatMasker. The SQuIRE call module was used to perform differential expression analysis of genes, or TE by family, or TE by sub-family between Control and Runx1$^{\Delta GMP}$ cells.

**Bulk RNA-seq**: Cells were sorted, and total RNA samples were isolated (RNeasy Mini Kit, Qiagen). Sample QC, library preparations, and sequencing reactions were conducted at GENEWIZ, LLC./Azenta US, Inc (South Plainfield, NJ, USA) as follows: RNA samples were quantified using Qubit 2.0 Fluorometer (ThermoFisher Scientific, Waltham, MA, USA) and RNA integrity was checked using TapeStation (Agilent Technologies). ERCC RNA Spike-In Mix kit (cat. 4456740) from ThermoFisher Scientific, was added to normalized cell number prior to library preparation following manufacturer's protocol. The RNA sequencing libraries were prepared using the NEBNext Ultra II RNA Library Prep Kit for Illumina using manufacturer's instructions (NEB). Briefly, mRNAs were initially enriched with Oligod(T) beads. Enriched mRNAs were fragmented for 15 min at 94 °C. First strand and second strand cDNA were subsequently synthesized. cDNA fragments were end repaired and adenylated at 3'ends, and universal adapters were ligated to cDNA fragments, followed by index addition and library enrichment by PCR with limited cycles. The sequencing libraries were validated on the Agilent TapeStation (Agilent Technologies), and quantified by using Qubit 2.0 Fluorometer (ThermoFisher Scientific) as well as by quantitative PCR (KAPA Biosystems). The sequencing libraries were multiplexed and clustered onto a flow cell. After clustering, the flowcell was loaded onto the NovaSeq 6000 instrument and sequenced using a 2x150bp Paired End (PE) configuration according to the manufacturer's instructions.

**Bulk RNA-seq data processing:** Sequencing and base calling was performed by GeneWiz (https://www.genewiz.com/). Raw sequence data (.bcl files) were converted into fastq files and de-multiplexed using Illumina Bcl2Fastq v2.20 software. One mis- match was allowed for index sequence identification. After investigating the quality of the raw data, sequence reads were trimmed to remove possible adapter sequences and nucleotides with poor quality using

Trimmomatic v.0.36. The trimmed reads were mapped to the *Mus musculus* reference genome available on ENSEMBL using the STAR aligner v.2.5.2b. BAM files were generated as a result of this step. Unique gene hit counts were calculated by using feature Counts from the Subread package v.1.5.2. Only unique reads that fell within exon regions were counted. After extraction of gene hit counts, the gene hit counts table was used for downstream differential expression analysis. Using DESeq2, a comparison of gene expression between the groups of samples was performed. The Wald test was used to generate P values and Log2 fold changes. Genes with adjusted P values $< 0.05$ and absolute log2 fold changes $>1$ were called as differentially expressed genes for each comparison. Merged replicates were used to create bigwig files for visualization using Deeptools v3.5.1 [60] with the following parameters: normalized to Bins Per Million mapped reads (BPM), bin size: 50

#### SUPPLEMENTARY FIGURES:

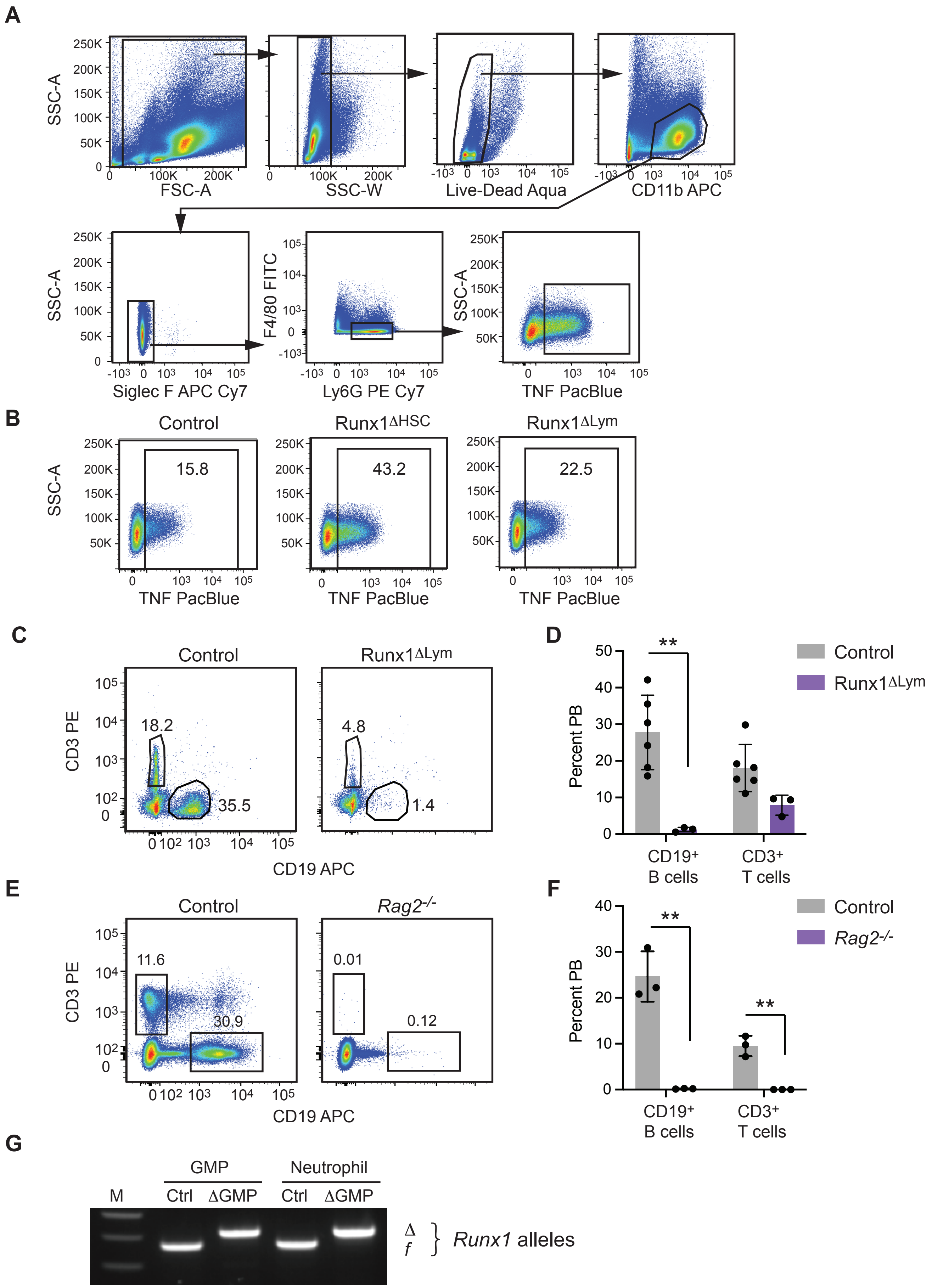
A
SSC-A
FSC-A
SSC-W
Live-Dead Aqua
CD11b APC
Siglec F APC Cy7
F4/80 FITC
Ly6G PE Cy7
TNF PacBlue
B
Control
Runx1ΔHSC
Runx1ΔLym
15.8
43.2
22.5
C
CD3 PE
CD19 APC
18.2
35.5
4.8
1.4
D
Percent PB
Control
Runx1ΔLym
CD19+ B cells
CD3+ T cells
E
Control
Rag2-/-
11.6
30.9
0.01
0.12
F
G
GMP
Neutrophil
M
Ctrl
ΔGMP
Δ
f
Runx1 alleles

**Supplementary Figure 1. RUNX1 function in GMPs is necessary to restrict inflammatory cytokine production by neutrophils. A)** Scatter plots showing gating strategy for neutrophil isolation and analysis. Shown is an analysis of $TNF^{+}$ $CD11b^{+}Ly6G^{+}SiglecF^{-}F4/80^{-}$ neutrophils**.B)** Scatter plots of intracellular TNF in Control, $Runx1^{\Delta HSC}$ and $Runx1^{\Delta Lym}$ neutrophils. **C)** Analysis of $CD19^{+}$ B cells and $CD3^{+}$ T cells in the PB of Control (C57BL6/J) and $Runx1^{\Delta Lym}$ mice in which *Runx1* floxed alleles were deleted with Rag1-Cre. **D)** Quantification of PB B and T cells in $Runx1^{\Delta Lym}$ mice. Mean ± SD, two-tailed, unpaired t-test, representative of 2 experiments, a total of 9 mice analyzed, **P≤0.01. **E)** Analysis of B and T cells in the PB of Control and *Rag2*$^{-/-}$ mice. **F)** Quantification of PB B and T cells in *Rag2*$^{-/-}$ mice, as in panel D. n= 6 mice analyzed. **G)** PCR showing *Runx1*$^{f/f}$ deletion with Cebpa-Cre in neutrophils and GMPs. Undeleted (f) and deleted (Δ) alleles are shown.

**Zezulin_S2**

**A**

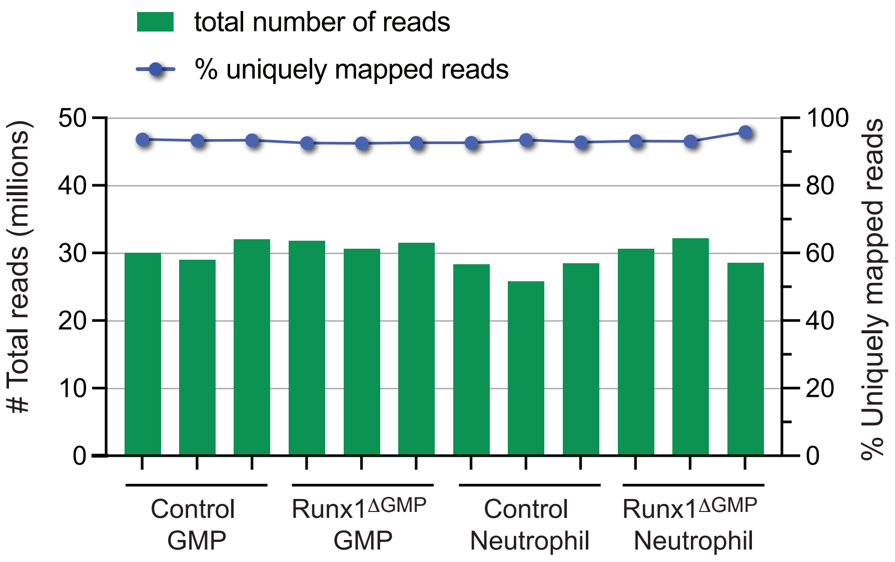


**B**

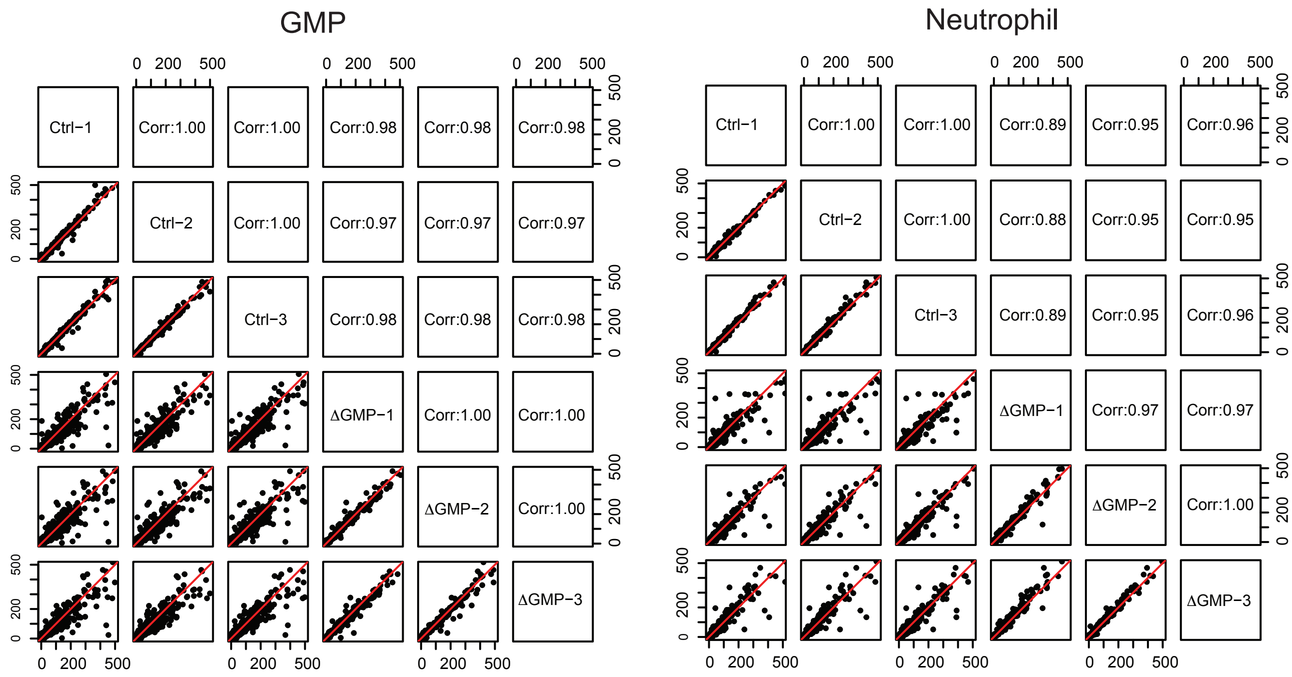

**Supplementary Figure 2. Sample sequencing statistics. A)** Mapping statistics of RNA-seq data. RNA-Seq reads were mapped to the *Mus musculus* GRCm38 ERCC reference genome available on ENSEMBL using the STAR aligner v.2.5.2b. Green bars show the number of total reads (in millions). Blue dots show the percentages of uniquely mapped reads. **B)** Pearson correlation of RNA-seq replicate samples. Scatter plot matrix of replicates. Pearson correlation coefficients were calculated based on transcripts per million (TPM) of genes for each pair of samples. The TPM of genes are plotted for each pair of samples, and the Pearson correlation coefficients are shown for each pair of samples.

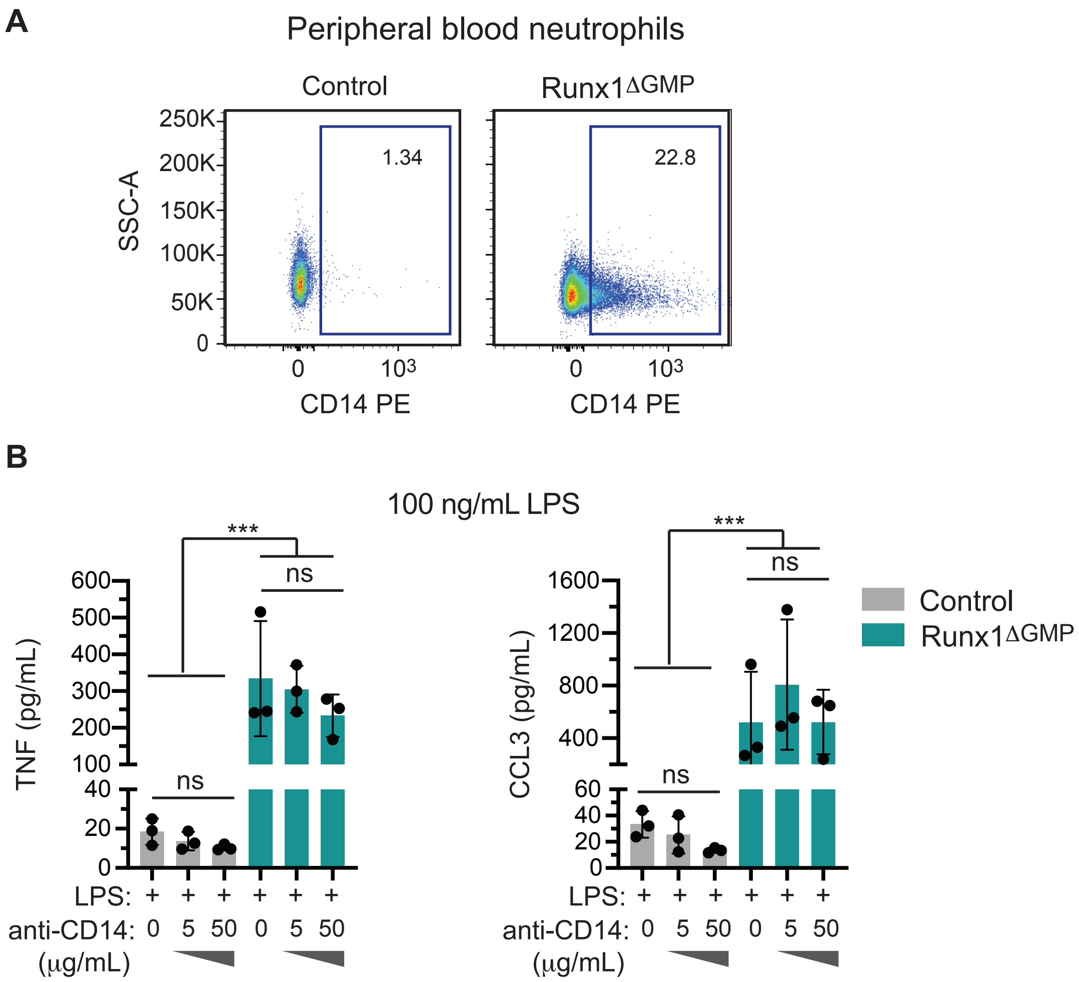
A
Peripheral blood neutrophils
Control
Runx1ΔGMP
SSC-A
250K
200K
150K
100K
50K
0
1.34
22.8
0
10³
CD14 PE
B
100 ng/mL LPS
***
ns
TNF (pg/mL)
600
500
400
300
200
100
40
20
0
CCL3 (pg/mL)
1600
1200
800
400
60
40
20
0
Control
Runx1ΔGMP
LPS: + + + + + +
anti-CD14: 0 5 50 0 5 50
(µg/mL)

**Supplementary Figure 3. RUNX1 loss results in elevated levels of key TLR4 signaling molecules. A)** Representative scatter plots of CD14 expression on PB neutrophils from Control and Runx1$^{\Delta GMP}$ mice. **B**) CBA analysis demonstrating effect of CD14 blocking antibody on TNF and CCL3 production by purified BM-derived neutrophils stimulated for 8 hours with vehicle or a high dose (100 ng/mL) of LPS. Mean ± SD, one-way ANOVA plus Tukey multiple comparison test, representative of 3 experiments, a total of 16 mice were analyzed. ***P=0.0001, ns= not significant.

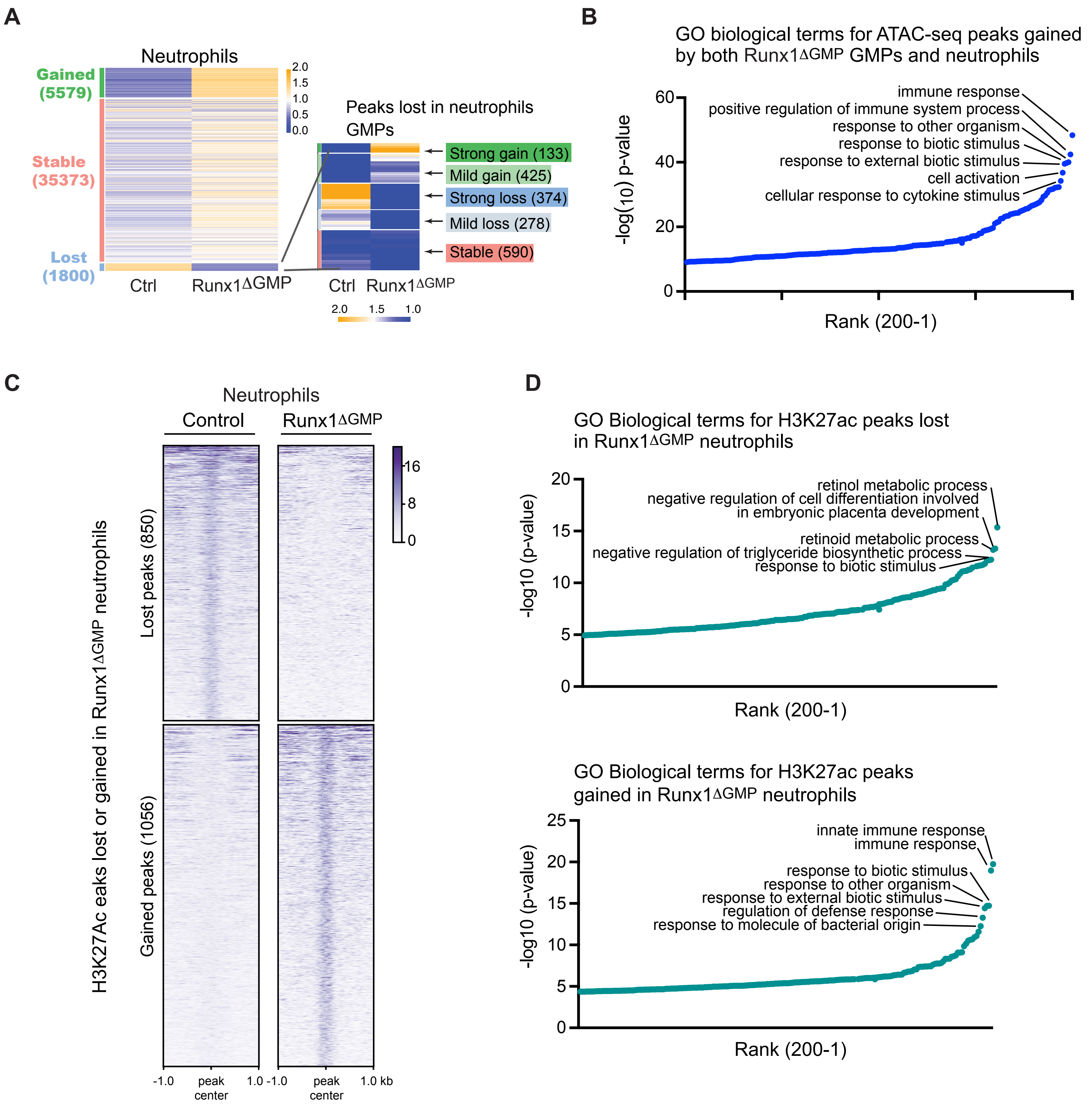
A
Neutrophils
Gained (5579)
Stable (35373)
Lost (1800)
Ctrl
Runx1ΔGMP
2.0
1.5
1.0
0.5
0.0
Peaks lost in neutrophils GMPs
Strong gain (133)
Mild gain (425)
Strong loss (374)
Mild loss (278)
Stable (590)
Ctrl
Runx1ΔGMP
2.0
1.5
1.0
B
GO biological terms for ATAC-seq peaks gained by both Runx1ΔGMP GMPs and neutrophils
immune response
positive regulation of immune system process
response to other organism
response to biotic stimulus
response to external biotic stimulus
cell activation
cellular response to cytokine stimulus
-log(10) p-value
Rank (200-1)
C
Neutrophils
Control
Runx1ΔGMP
H3K27Ac eaks lost or gained in Runx1ΔGMP neutrophils
Lost peaks (850)
Gained peaks (1056)
16
8
0
-1.0
peak center
1.0 -1.0
peak center
1.0 kb
D
GO Biological terms for H3K27ac peaks lost in Runx1ΔGMP neutrophils
retinol metabolic process
negative regulation of cell differentiation involved in embryonic placenta development
retinoid metabolic process
negative regulation of triglyceride biosynthetic process
response to biotic stimulus
-log10 (p-value)
Rank (200-1)
GO Biological terms for H3K27ac peaks gained in Runx1ΔGMP neutrophils
innate immune response
immune response
response to biotic stimulus
response to other organism
response to external biotic stimulus
regulation of defense response
response to molecule of bacterial origin
-log10 (p-value)
Rank (200-1)

**Supplementary Figure 4. Loss of RUNX1 activates genes that mediate innate immune responses. A)** Left, heatmap of ATAC-seq signals for Control (Ctrl) and Runx1$^{\Delta GMP}$ neutrophils. Peaks are categorized into Gained, Lost (in Runx1$^{\Delta GMP}$ GMPs), and Stable groups. Right, heatmap of ATAC-seq signal in Control and Runx1$^{\Delta GMP}$ GMPs for peaks lost in Runx1$^{\Delta GMP}$ neutrophils. **B)** GO biological terms for gained ATAC-seq peaks shared by Runx1$^{\Delta GMP}$ GMPs and neutrophils. **C)** Heat maps of H3K27ac signals in Control and Runx1$^{\Delta GMP}$ neutrophils (scales are normalized to RPGC read counts) for 850 lost and 1056 gained regions following RUNX1 loss. **D)** Gene Ontology (GO) analysis of genes with H3K27ac lost or gained peaks in Control and Runx1$^{\Delta GMP}$ neutrophils. The top 200 GO terms are plotted.

**Zezulin_S5**

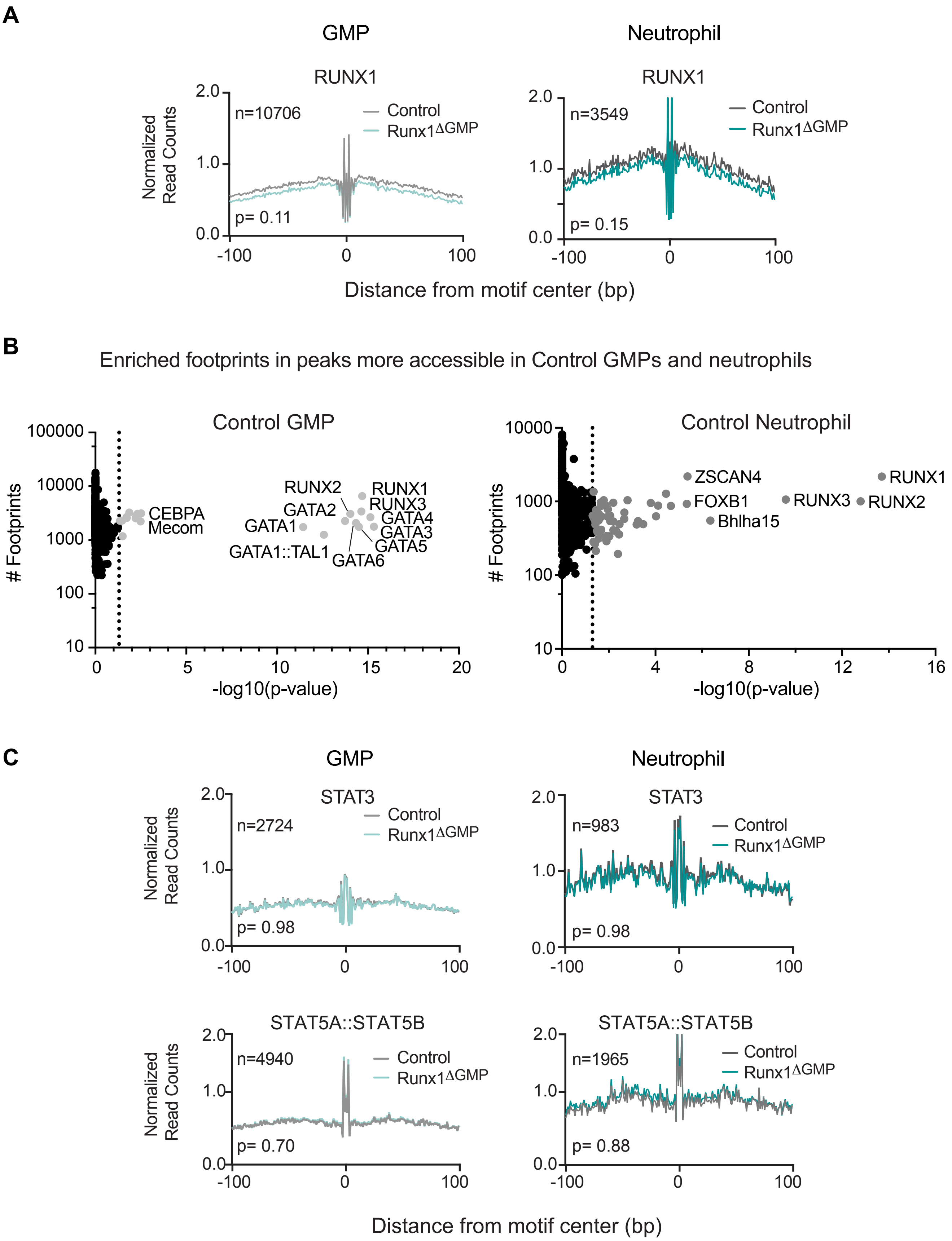
A
GMP
Neutrophil
RUNX1
RUNX1
Normalized Read Counts
n=10706
Control
Runx1ΔGMP
p= 0.11
n=3549
Control
Runx1ΔGMP
p= 0.15
2.0
1.0
0.0
-100
0
100
Distance from motif center (bp)
B
Enriched footprints in peaks more accessible in Control GMPs and neutrophils
Control GMP
Control Neutrophil
### Footprints
100000
10000
1000
100
10
0
5
10
15
20
-log10(p-value)
CEBPA
Mecom
RUNX2
RUNX1
RUNX3
GATA2
GATA4
GATA1
GATA3
GATA5
GATA1::TAL1
GATA6
ZSCAN4
FOXB1
Bhlha15
RUNX3
RUNX1
RUNX2
4
8
12
16
C
GMP
Neutrophil
STAT3
STAT3
n=2724
p= 0.98
n=983
p= 0.98
STAT5A::STAT5B
STAT5A::STAT5B
n=4940
p= 0.70
n=1965
p= 0.88
Control
Runx1ΔGMP
Normalized Read Counts
Distance from motif center (bp)

**Supplementary Figure 5. Loss of RUNX1 affects type I IFN signaling. A)** Footprint profile plots for RUNX1 showing average normalized read counts and p-values calculated by HINT-differential [29] using all peaks in Control and Runx1$^{\Delta GMP}$ GMPs and neutrophils. **B)** Scatter plots showing enriched TF footprints in regions of chromatin with decreased accessibility in Runx1$^{\Delta GMP}$ neutrophils and GMPs relative to Controls. The number of footprints for each TF at regions of chromatin with decreased accessibility in Runx1$^{\Delta GMP}$ neutrophils and GMPs is displayed on the y-axis for the Control cells. Colored circles indicate $p<0.05$; p-value calculated using biFET. **C)** Digital footprint profile plots for selected STAT TFs showing average normalized read counts and p-values calculated with HINT-differential using all peaks in Control and Runx1$^{\Delta GMP}$ cells.

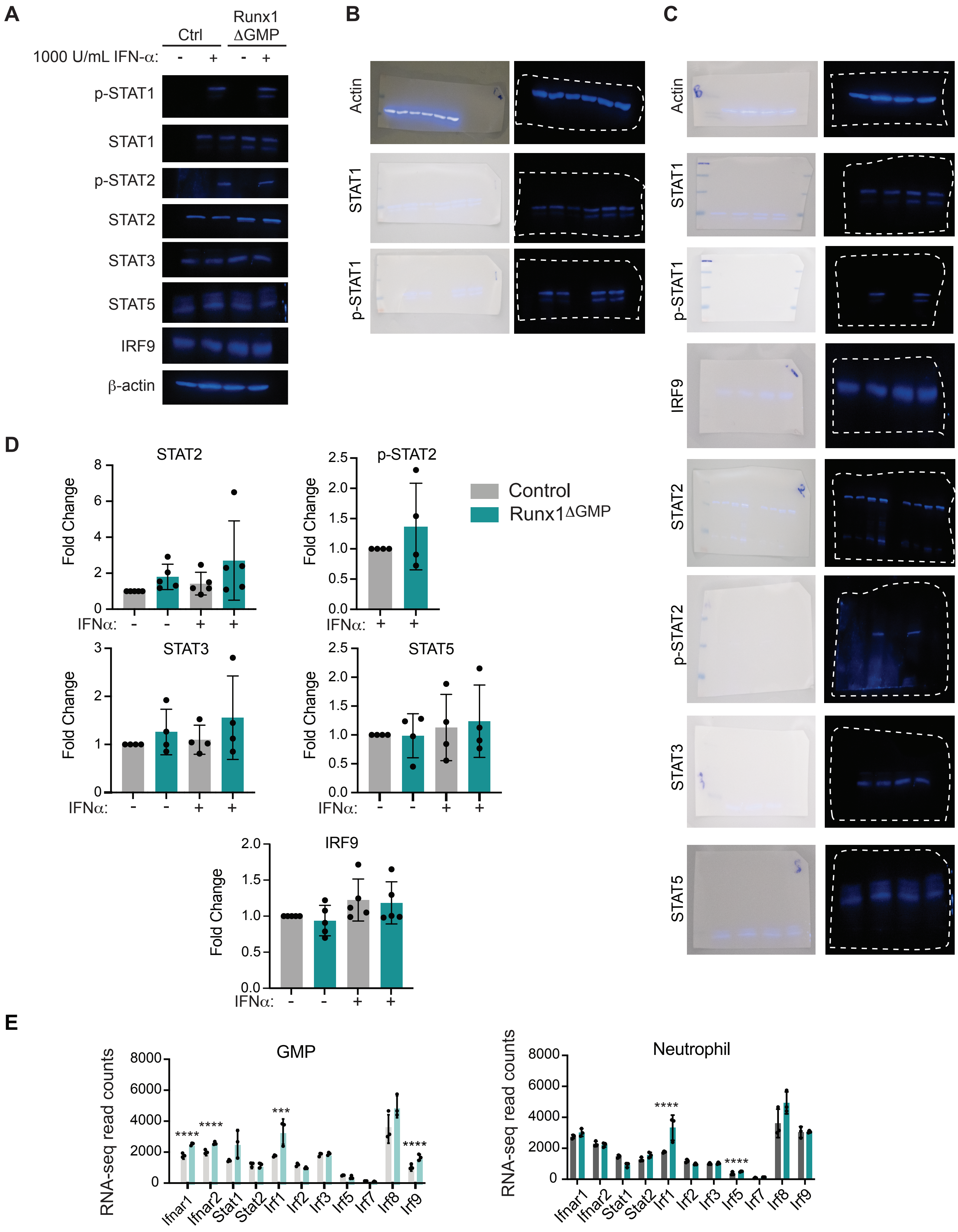
A
B
C
D
E
Ctrl
Runx1 ΔGMP
1000 U/mL IFN-α:
p-STAT1
STAT1
p-STAT2
STAT2
STAT3
STAT5
IRF9
β-actin
Actin
Control
Runx1ΔGMP
Fold Change
IFNα:
RNA-seq read counts
GMP
Neutrophil
Ifnar1
Ifnar2
Stat1
Stat2
Irf1
Irf2
Irf3
Irf5
Irf7
Irf8
Irf9

**Supplementary Figure 6. Loss of RUNX1 increases the levels of a subset of type I IFN signaling molecules. A)** Western blot for total STAT, phosphorylated STAT (p-STAT), and IRF9 plus a β-actin control in Control and Runx1$^{\Delta GMP}$ neutrophils in the presence or absence of IFN-α. B. **B)** Original Western blots for STAT, p-STAT, IRF, and β-actin control in Control and Runx1$^{\Delta GMP}$ neutrophils in Figure 4D. The edges of each blot are outlined with a dotted line. **C)** Original Western blots for STAT, pSTAT, IRF, and β-actin control in Control and Runx1$^{\Delta GMP}$ neutrophils in panel A. **D)** Quantification of Western blots for STAT2, p-STAT2, STAT2, STAT5, and IRF9. ANOVA plus Tukey multiple comparison tests; all differences except for STAT1 and p-STAT1 in Fig. 4D were not significant. **E)** RNA-seq read counts for genes encoding several components of the type I IFN signaling pathway including receptors (*Ifnar1, Ifnar2*), STATs, and IRFs in GMPs and neutrophils. Mean ± SD, unpaired two-tailed t test. ***P=0.0001, ****P<0.0001.

**Zezulin_S7**

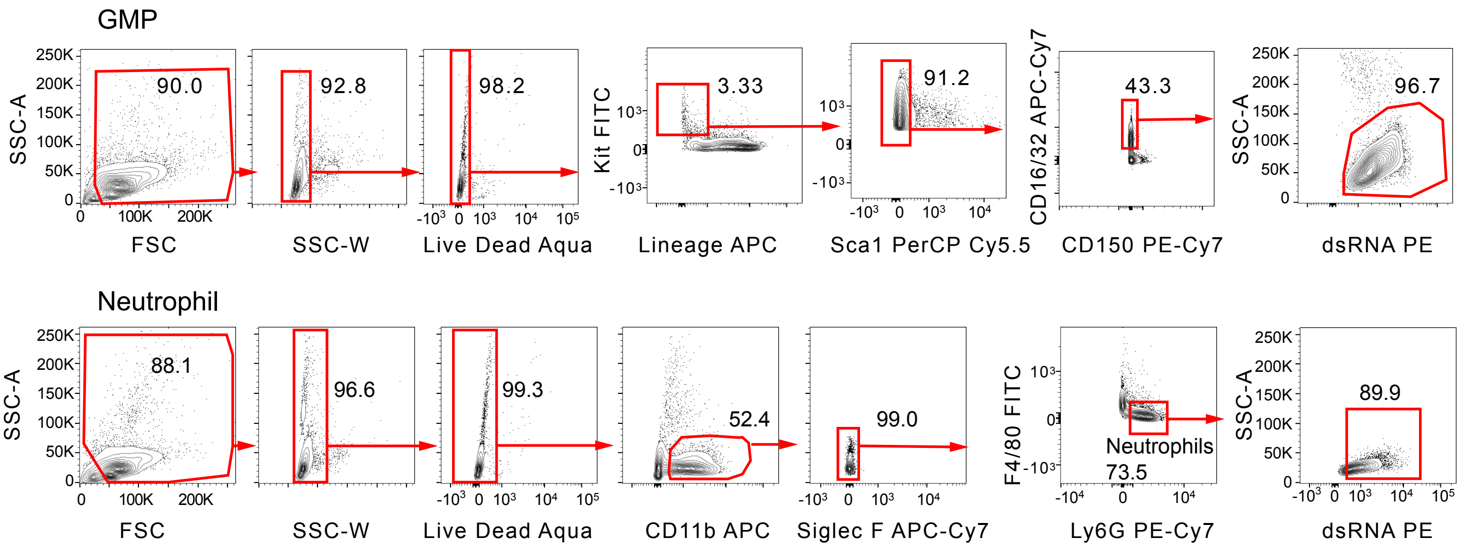
GMP
SSC-A
FSC
90.0
SSC-W
92.8
Live Dead Aqua
98.2
Kit FITC
Lineage APC
3.33
Sca1 PerCP Cy5.5
91.2
CD16/32 APC-Cy7
CD150 PE-Cy7
43.3
dsRNA PE
96.7
Neutrophil
88.1
96.6
99.3
CD11b APC
52.4
Siglec F APC-Cy7
99.0
F4/80 FITC
Ly6G PE-Cy7
Neutrophils
73.5
89.9

**Supplementary Figure 7. Gating strategy for analyzing dsRNA in GMPs and neutrophils.** Quantile contour FACS plots depicting gating strategies. Numbers on the x and y-axes are indicated on the first plot on the left, and unless changed are not depicted on plots to the right of the preceding plot.

**Supplementary Table 1:** List of all antibodies used for flow cytometry, western blot analysis, and H3K27ac ChIP-seq. For each antibody, the clone, fluorophore, dilution, manufacturer, and antibody registry number (or manufacturer's catalog number when no RRID number is available) are provided.

**Supplementary Table 2**: GO terms for peaks gained in Runx1$^{\Delta GMP}$ GMPs and neutrophils.

**Supplementary Table 3:** Table showing differential activity of transcription factors calculated using HINT-differential and enriched transcription factor footprints at regions of chromatin with increased accessibility calculated using BiFET v.1.16.0 software.

**Supplementary Table 4:** TE subfamilies that were enriched or depleted in the immunoprecipitate of Runx1$^{\Delta GMP}$ neutrophils.

1 Bellissimo, D. C. & Speck, N. A. RUNX1 Mutations in Inherited and Sporadic Leukemia. *Front Cell Dev Biol* **5**, 111, doi:10.3389/fcell.2017.00111 (2017).
2 Deuitch, N., Broadbridge, E., Cunningham, L. & Liu, P. P. RUNX1 Familial Platelet Disorder with Associated Myeloid Malignancies. *Gene Reviews* (2021).
3 Brown, A. L. *et al.* RUNX1-mutated families show phenotype heterogeneity and a somatic mutation profile unique to germline predisposed AML. *Blood Adv* **4**, 1131-1144, doi:10.1182/bloodadvances.2019000901 (2020).
4 Churpek, J. E. *et al.* Genomic analysis of germ line and somatic variants in familial myelodysplasia/acute myeloid leukemia. *Blood* **126**, 2484-2490, doi:10.1182/blood-2015-04-641100 (2015).
5 Sorrell, A. *et al.* Hereditary leukemia due to rare RUNX1c splice variant (L472X) presents with eczematous phenotype. *Int J Clin Med* **3**, doi:10.4236/ijcm.2012.37110 (2012).
6 Sacco, K. *et al.* Germline RUNX1 deficiency predisposes to allergy and autoimmunity. *The Journal of Allergy and Clinical Immunology* **147**, doi:https://doi.org/10.1016/j.jaci.2020.12.266 (2020).
7 Kouroukli, O., Symeonidis, A., Foukas, P., Maragkou, M. K. & Kourea, E. P. Bone Marrow Immune Microenvironment in Myelodysplastic Syndromes. *Cancers (Basel)* **14**, doi:10.3390/cancers14225656 (2022).
8 Stubbins, R. J., Platzbecker, U. & Karsan, A. Inflammation and myeloid malignancy: quenching the flame. *Blood* **140**, 1067-1074, doi:10.1182/blood.2021015162 (2022).
9 Luo, M. C. *et al.* Runt-related Transcription Factor 1 (RUNX1) Binds to p50 in Macrophages and Enhances TLR4-triggered Inflammation and Septic Shock. *The Journal of biological chemistry* **291**, 22011-22020, doi:10.1074/jbc.M116.715953 (2016).
10 Ono, M. *et al.* Foxp3 controls regulatory T-cell function by interacting with AML1/Runx1. *Nature* **446**, 685-689 (2007).
11 Tang, X. *et al.* Runt-Related Transcription Factor 1 Regulates LPS-Induced Acute Lung Injury via NF-κB Signaling. *American Journal of Respiratory Cell and Molecular Biology* **57**, 174-183, doi:10.1165/rcmb.2016-0319OC (2017).
12 DeKelver, R. C. *et al.* RUNX1-ETO induces a type I interferon response which negatively effects t(8;21)-induced increased self-renewal and leukemia development. *Leuk Lymphoma* **55**, 884-891, doi:10.3109/10428194.2013.815351 (2014).
13 Hu, Y. *et al.* RUNX1 inhibits the antiviral immune response against influenza A virus through attenuating type I interferon signaling. *Virol J* **19**, 39, doi:10.1186/s12985-022-01764-8 (2022).
14 Thomsen, I. *et al.* RUNX1 Regulates a Transcription Program That Affects the Dynamics of Cell Cycle Entry of Naive Resting B Cells. *J Immunol* **207**, 2976-2991, doi:10.4049/jimmunol.2001367 (2021).
15 Geis, F. K. & Goff, S. P. Silencing and Transcriptional Regulation of Endogenous Retroviruses: An Overview. *Viruses* **12**, doi:10.3390/v12080884 (2020).
16 Matteucci, C., Balestrieri, E., Argaw-Denboba, A. & Sinibaldi-Vallebona, P. Human endogenous retroviruses role in cancer cell stemness. *Semin Cancer Biol* **53**, 17-30, doi:10.1016/j.semcancer.2018.10.001 (2018).
17 Kitsou, K., Lagiou, P. & Magiorkinis, G. Human Endogenous Retroviruses in Cancer: Oncogenesis mechanisms and Clinical Implications. *J Med Virol*, doi:10.1002/jmv.28350 (2022).
18 Saleh, A., Macia, A. & Muotri, A. R. Transposable Elements, Inflammation, and Neurological Disease. *Front Neurol* **10**, 894, doi:10.3389/fneur.2019.00894 (2019).

19 Chuong, E. B., Elde, N. C. & Feschotte, C. Regulatory evolution of innate immunity through co-option of endogenous retroviruses. *Science* **351**, 1083-1087, doi:10.1126/science.aad5497 (2016).

20 Chen, Y. G. & Hur, S. Cellular origins of dsRNA, their recognition and consequences. *Nat Rev Mol Cell Biol* **23**, 286-301, doi:10.1038/s41580-021-00430-1 (2022).

21 Colombo, A. R., Triche, T., Jr. & Ramsingh, G. Transposable Element Expression in Acute Myeloid Leukemia Transcriptome and Prognosis. *Sci Rep* **8**, 16449, doi:10.1038/s41598-018-34189-x (2018).

22 Bellissimo, D. C. *et al.* Runx1 negatively regulates inflammatory cytokine production by neutrophils in response to Toll-like receptor signaling. *Blood Adv* **4**, 1145-1158, doi:10.1182/bloodadvances.2019000785 (2020).

23 Shinkai, Y. *et al.* RAG-2-deficient mice lack mature lymphocytes owing to inability to initiate V(D)J rearrangement. *Cell* **68**, 855-867 (1992).

24 Abram, C. L., Roberge, G. L., Hu, Y. & Lowell, C. A. Comparative analysis of the efficiency and specificity of myeloid-Cre deleting strains using ROSA-EYFP reporter mice. *Journal of immunological methods* **408**, 89-100, doi:10.1016/j.jim.2014.05.009 (2014).

25 Zanoni, I. & Granucci, F. Role of CD14 in host protection against infections and in metabolism regulation. *Front Cell Infect Microbiol* **3**, 32, doi:10.3389/fcimb.2013.00032 (2013).

26 Ciesielska, A., Matyjek, M. & Kwiatkowska, K. TLR4 and CD14 trafficking and its influence on LPS-induced pro-inflammatory signaling. *Cell Mol Life Sci* **78**, 1233-1261, doi:10.1007/s00018-020-03656-y (2021).

27 Moore, K. J. *et al.* Divergent response to LPS and bacteria in CD14-deficient murine macrophages. *J Immunol* **165**, 4272-4280, doi:10.4049/jimmunol.165.8.4272 (2000).

28 Perera, P. Y., Vogel, S. N., Detore, G. R., Haziot, A. & Goyert, S. M. CD14-dependent and CD14-independent signaling pathways in murine macrophages from normal and CD14 knockout mice stimulated with lipopolysaccharide or taxol. *J Immunol* **158**, 4422-4429 (1997).

29 Li, Z. *et al.* Identification of transcription factor binding sites using ATAC-seq. *Genome Biol* **20**, 45, doi:10.1186/s13059-019-1642-2 (2019).

30 Youn, A., Marquez, E. J., Lawlor, N., Stitzel, M. L. & Ucar, D. BiFET: sequencing Bias-free transcription factor Footprint Enrichment Test. *Nucleic Acids Res* **47**, e11, doi:10.1093/nar/gky1117 (2019).

31 Espinet, E. *et al.* Aggressive PDACs Show Hypomethylation of Repetitive Elements and the Execution of an Intrinsic IFN Program Linked to a Ductal Cell of Origin. *Cancer Discov* **11**, 638-659, doi:10.1158/2159-8290.CD-20-1202 (2021).

32 Wang, W. *et al.* Unphosphorylated ISGF3 drives constitutive expression of interferon-stimulated genes to protect against viral infections. *Sci Signal* **10**, doi:10.1126/scisignal.aah4248 (2017).

33 Platanitis, E. *et al.* A molecular switch from STAT2-IRF9 to ISGF3 underlies interferon-induced gene transcription. *Nature communications* **10**, 2921, doi:10.1038/s41467-019-10970-y (2019).

34 Gazquez-Gutierrez, A., Witteveldt, J., S, R. H. & Macias, S. Sensing of transposable elements by the antiviral innate immune system. *RNA* **27**, 735-752, doi:10.1261/rna.078721.121 (2021).

35 Son, K. N., Liang, Z. & Lipton, H. L. Double-Stranded RNA Is Detected by Immunofluorescence Analysis in RNA and DNA Virus Infections, Including Those by Negative-Stranded RNA Viruses. *J Virol* **89**, 9383-9392, doi:10.1128/JVI.01299-15 (2015).

36 Song, W.-J. *et al.* Haploinsufficiency of *CBFA2* (*AML1*) causes familial thrombocytopenia with propensity to develop acute myelogenous leukamia (FPD/AML). *Nature Genet.* **23**, 166-175 (1999).
37 Homan, C. C. *et al.* The RUNX1 database (RUNX1db): establishment of an expert curated RUNX1 registry and genomics database as a public resource for familial platelet disorder with myeloid malignancy. *Haematologica* **106**, 3004-3007, doi:10.3324/haematol.2021.278762 (2021).
38 Kanno, T. *et al.* Intrinsic transcriptional activation-inhibition domains of the polyomavirus enhancer binding protein 2/core binding factor a subunit revealed in the presence of the b subunit. *Mol Cell Biol.* **18**, 2444-2454 (1998).
39 Kalafati, L., Hatzioannou, A., Hajishengallis, G. & Chavakis, T. The role of neutrophils in trained immunity. *Immunol Rev*, doi:10.1111/imr.13142 (2022).
40 Christ, A. *et al.* Western Diet Triggers NLRP3-Dependent Innate Immune Reprogramming. *Cell* **172**, 162-175 e114, doi:10.1016/j.cell.2017.12.013 (2018).
41 Li, X. *et al.* Maladaptive innate immune training of myelopoiesis links inflammatory comorbidities. *Cell* **185**, 1709-1727 e1718, doi:10.1016/j.cell.2022.03.043 (2022).
42 Kalafati, L. *et al.* Innate Immune Training of Granulopoiesis Promotes Anti-tumor Activity. *Cell* **183**, 771-785 e712, doi:10.1016/j.cell.2020.09.058 (2020).
43 Moorlag, S. *et al.* BCG Vaccination Induces Long-Term Functional Reprogramming of Human Neutrophils. *Cell reports* **33**, 108387, doi:10.1016/j.celrep.2020.108387 (2020).
44 Hosokawa, H. *et al.* Bcl11b sets pro-T cell fate by site-specific cofactor recruitment and by repressing Id2 and Zbtb16. *Nat Immunol* **19**, 1427-1440, doi:10.1038/s41590-018-0238-4 (2018).
45 Manghera, M. & Douville, R. N. Endogenous retrovirus-K promoter: a landing strip for inflammatory transcription factors? *Retrovirology* **10**, 16, doi:10.1186/1742-4690-10-16 (2013).
46 Jaiswal, S. & Libby, P. Clonal haematopoiesis: connecting ageing and inflammation in cardiovascular disease. *Nat Rev Cardiol* **17**, 137-144, doi:10.1038/s41569-019-0247-5 (2020).
47 Jaiswal, S. *et al.* Age-related clonal hematopoiesis associated with adverse outcomes. *N Engl J Med* **371**, 2488-2498, doi:10.1056/NEJMoa1408617 (2014).
48 Genovese, G. *et al.* Clonal hematopoiesis and blood-cancer risk inferred from blood DNA sequence. *N Engl J Med* **371**, 2477-2487, doi:10.1056/NEJMoa1409405 (2014).
49 Growney, J. D. *et al.* Loss of Runx1 perturbs adult hematopoiesis and is associated with a myeloproliferative phenotype. *Blood* **106**, 494-504 (2005).
50 Stadtfeld, M. & Graf, T. Assessing the role of hematopoietic plasticity for endothelial and hepatocyte development by non-invasive lineage tracing. *Development* **132**, 203-213 (2005).
51 McCormack, M. P., Forster, A., Drynan, L., Pannell, R. & Rabbitts, T. H. The LMO2 T-cell oncogene is activated via chromosomal translocations or retroviral insertion during gene therapy but has no mandatory role in normal T-cell development. *Mol Cell Biol* **23**, 9003-9013, doi:10.1128/MCB.23.24.9003-9013.2003 (2003).
52 Wolfler, A. *et al.* Lineage-instructive function of C/EBPalpha in multipotent hematopoietic cells and early thymic progenitors. *Blood* **116**, 4116-4125, doi:10.1182/blood-2010-03-275404 (2010).
53 Edgar, R., Domrachev, M. & Lash, A. E. Gene Expression Omnibus: NCBI gene expression and hybridization array data repository. *Nucleic Acids Res* **30**, 207-210, doi:10.1093/nar/30.1.207 (2002).
54 Kagoshima, H. *et al.* The Runt-domain identifies a new family of heteromeric DNA-binding transcriptional regulatory proteins. *Trends Genet.* **9**, 338-341 (1993).

55 Langmead, B. & Salzberg, S. L. Fast gapped-read alignment with Bowtie 2. *Nat Methods* **9**, 357-359, doi:10.1038/nmeth.1923 (2012).
56 Langmead, B., Trapnell, C., Pop, M. & Salzberg, S. L. Ultrafast and memory-efficient alignment of short DNA sequences to the human genome. *Genome Biol* **10**, R25, doi:gb-2009-10-3-r25 [pii]
10.1186/gb-2009-10-3-r25 (2009).
57 Danecek, P. *et al.* Twelve years of SAMtools and BCFtools. *Gigascience* **10**, doi:10.1093/gigascience/giab008 (2021).
58 Tarasov, A., Vilella, A. J., Cuppen, E., Nijman, I. J. & Prins, P. Sambamba: fast processing of NGS alignment formats. *Bioinformatics* **31**, 2032-2034, doi:10.1093/bioinformatics/btv098 (2015).
59 Zhang, Y. *et al.* Model-based analysis of ChIP-Seq (MACS). *Genome Biol* **9**, R137, doi:gb-2008-9-9-r137 [pii]
10.1186/gb-2008-9-9-r137 (2008).
60 Ramirez, F. *et al.* deepTools2: a next generation web server for deep-sequencing data analysis. *Nucleic Acids Res* **44**, W160-165, doi:10.1093/nar/gkw257 (2016).
61 Hiller, M. *et al.* Computational methods to detect conserved non-genic elements in phylogenetically isolated genomes: application to zebrafish. *Nucleic Acids Res* **41**, e151, doi:10.1093/nar/gkt557 (2013).
62 McLean, C. Y. *et al.* GREAT improves functional interpretation of cis-regulatory regions. *Nat Biotechnol* **28**, 495-501, doi:nbt.1630 [pii]
10.1038/nbt.1630 (2010).
63 Buenrostro, J. D., Giresi, P. G., Zaba, L. C., Chang, H. Y. & Greenleaf, W. J. Transposition of native chromatin for fast and sensitive epigenomic profiling of open chromatin, DNA-binding proteins and nucleosome position. *Nat Methods* **10**, 1213-1218, doi:10.1038/nmeth.2688 (2013).
64 Martin, M. Cutadapt Removes Adapter Sequences From High-Throughput Sequencing Reads. *EMBnet.journal* **17** (2011).
65 Quinlan, A. R. & Hall, I. M. BEDTools: a flexible suite of utilities for comparing genomic features. *Bioinformatics* **26**, 841-842, doi:10.1093/bioinformatics/btq033 (2010).
66 Heinz, S. *et al.* Simple combinations of lineage-determining transcription factors prime cis-regulatory elements required for macrophage and B cell identities. *Mol Cell* **38**, 576-589, doi:10.1016/j.molcel.2010.05.004 (2010).
67 Gusmao, E. G., Allhoff, M., Zenke, M. & Costa, I. G. Analysis of computational footprinting methods for DNase sequencing experiments. *Nat Methods* **13**, 303-309, doi:10.1038/nmeth.3772 (2016).
68 Tavora, B. *et al.* Tumoural activation of TLR3-SLIT2 axis in endothelium drives metastasis. *Nature* **586**, 299-304, doi:10.1038/s41586-020-2774-y (2020).
69 Yang, W. R., Ardeljan, D., Pacyna, C. N., Payer, L. M. & Burns, K. H. SQuIRE reveals locus-specific regulation of interspersed repeat expression. *Nucleic Acids Res* **47**, e27, doi:10.1093/nar/gky1301 (2019).
